# Random and non-random variation in flower color along an urban-rural gradient in the introduced mustard *Hesperis matronalis*

**DOI:** 10.1101/2025.04.22.650098

**Authors:** Katherine G. Maunder, Kaitlyn Dawson, Lauryn Joslin, Lucas Eckert, Marlene M. Kraml, Chloë Dean-Moore, Christopher G. Eckert

**Affiliations:** Department of Biology, Queen’s University, Kingston, Ontario K7L 3N6 Canada; Department of Ecology and Evolutionary Biology, University of Toronto, Toronto, Ontario M5S 3B2 Canada; Department of Biology, McGill University, Montréal, Quebec H3A 1B1 Canada

**Author notes:** **Correspondence**: Katherine G. Maunder, Department of Ecology and Evolutionary Biology, University of Toronto, Toronto, Ontario M5S 3B2 Canada.

**Keywords:** Brassicaceae, flower color, genetic drift, *Hesperis matronalis*, natural selection, nonindigenous plants, polymorphism, seed predation

## Abstract

**Premise:** Urbanization can alter the interplay of stochastic genetic drift and natural selection but these effects will depend on the biology and history of a species. To explore the influences of drift and selection we investigated patterns of flower color variation among populations of the introduced ornamental mustard, *Hesperis matronalis,* along an urban-rural gradient in eastern Ontario Canada.

**Methods:** We surveyed 136 naturalized stands of *H. matronalis* over three generations, and for each stand estimated the diversity of the three color morphs (white, pink, purple), the number of reproductive plants, and the degree of urbanization based on night sky brightness.

**Key Results:** Flower color morph diversity increased with both stand size and urbanization which is consistent with effects of genetic drift during colonization combined with multiple introductions of this horticultural plant in urban areas. However, the frequency and fixation of the purple morph systematically increased towards the rural end of the gradient. Although lifetime seed production did not vary among morphs, pre-dispersal seed predation by a recently adventive weevil was higher in the purple morph, particularly in rural areas. Estimated seed production in the absence of predation suggests a previous fitness advantage for the purple and pink morphs in rural areas and for the white morph in urban areas.

**Conclusions:** Random variation in flower color diversity may be influenced by stochastic processes and colonization history, while systematic variation in color morph frequencies may reflect past fitness differences among morphs that have been recently erased by seed predation.

## INTRODUCTION

Human activity is changing the evolutionary dynamics of numerous species, especially in urban habitats (Johnson and Munshi-South, 2017; Alberti et al. 2020). Urbanization can expose species to novel environments and may alter the form and strength of selection. This may create clines in ecologically important traits along urban-rural gradients (e.g. Thompson et al., 2016; Santangelo et al., 2022). However, urban areas can also differ from surrounding rural areas in the quantity, structure, and isolation of appropriate habitats (Miles et al., 2019), which may alter neutral processes such as genetic drift and gene flow. While it can be challenging to identify the relative contributions of selective and neutral processes (Santangelo et al., 2018), cities provide the opportunity to study the dynamic interplay of these evolutionary forces in novel widely available ecosystems. Beyond adding to our understanding of fundamental concepts, teasing apart the eco-evolutionary consequences of urbanization is also critical for effective urban planning and conservation efforts (Alberti et al., 2020).

A rich body of population genetic theory suggests that while all major evolutionary processes can be influenced by urbanization, the direction and magnitude of these effects will depend on the ecology of a particular species and hence how that species responds to urbanization (Miles et al. 2019). For many native species, urbanization fragments appropriate habitat and potentially reduces the genetic variation within urban populations. In this case, fragmentation is expected to a) decrease population size thereby increasing the loss of variation due to genetic drift and b) increase the spatial isolation of populations thereby restricting the replenishment of variation via gene flow (Noël and Lapointe 2010; Bartlewicz et al. 2015; Munshi-South et al. 2016; Rivkin and Johnson 2022). However, the opposite pattern may be observed in species that are well suited to urban environments or that are dispersed by human activity (Cadotte et al. 2017; Miles et al. 2019). For example, non-native plants have a long history of being introduced to urban areas through horticulture (Reichard and White 2001; Mack and Erneberg 2002; Harris et al. 2009; van Kleunen et al. 2018) and their subsequent range expansion after introduction is often facilitated by human activity (Kowarik 2003; Dehnen-Schmutz et al. 2007; McLean et al. 2017). In such cases, populations may be larger and less spatially isolated in urban areas compared to rural areas and thus better able to maintain genetic variation in the face of genetic drift. In addition, any stochastic reduction in genetic variation associated with founder effect during individual introductions can be offset by multiple introductions of a species (Barney 2006; Pairon et al. 2010; Roman and Darling 2007; Dlugosch and Parker 2008; Vicente et al. 2021) which may be particularly common in urban areas (Lu et al. 2022; Mairal et al. 2022). Thus, urban populations of introduced horticultural plants can serve as genetically diverse launching sites for the invasion of surrounding rural areas. During subsequent range expansion, sequential founder effect will likely decrease genetic diversity in populations further from the introduction sites (Austerlitz et al. 1997; Slatkin and Excoffier 2012) thereby creating a pattern of declining population genetic diversity from urban to rural areas. In this way, clines in phenotypic and genetic diversity along urban-rural gradients may be the result of neutral, non-selective forces.

Urbanization can also change the abiotic and biotic environment that a species experiences which may lead to novel selective pressures (Johnson and Munshi-South 2017). Altered selection may be indicated if the spatial pattern of phenotypic variation along an urban-rural gradient differs from the patterns expected from the neutral processes of genetic drift and gene flow alone. For instance, in a species whose dispersal is assisted by human activity, gene flow should be reduced and drift increased in more rural populations. As a result of these neutral processes, genetic and phenotypic variation should be reduced in rural areas and this reduction should be random with no bias towards any particular allele or phenotype (Hedrick 2000; but see Santangelo et al. 2018). In contrast, systematic changes in allele or phenotype frequencies are suggestive of variation in selection along the urban-rural gradient. For example, consistently high frequencies of wing-less queen morphs in the ant species *Myrmecina graminicola*ants in areas of high urbanization and habitat fragmentation suggests selection for lower dispersal abilities in more urban areas (Finand et al. 2023). Discrete phenotypic polymorphisms have provided excellent opportunities to study the interplay of neutral and selective forces in natural populations (Schemske and Bierzychudek 2007; Barrett 2019). If a polymorphism has a relatively simple genetic basis, then population variation in discrete phenotypic “morphs” may reflect genetic variation at the underlying trait loci (Joron et al. 2006; Kingsley et al. 2009)

In this study, we investigate the role of stochastic and selective processes affecting flower color variation in *Hesperis matronalis* L. (Brassicaceae, dame’s rocket). Studies of flower color variation were central to the debate over the prevalence of neutral versus selective processes influencing phenotypic variation in natural populations (Epling and Dobzhansky 1942; Schemske and Bierzychudek 2001). Yet since then, few studies have investigated the processes influencing the pattern of color variation across space or time (Sapir et al. 2021). Flower color is often thought to be under selection (Faegri and van der Pijl 1979), although definitive evidence of this is usually lacking for most species studied (Rausher 2008). Nevertheless, selection is often suspected because flower color can influence the foraging behaviours of pollinators which can lead to changes in gene transmission and ultimately individual fitness (Waser and Price 1981; Meléndez-Ackerman and Campbell 1998). However, flower color can also influence how plants respond to a wide variety of biotic and abiotic factors (Strauss and Whittall 2007; Caruso et al. 2018; Koski and Galloway 2020). Many of the pigments responsible for petal color, such as anthocyanins, also play a role in stress tolerance and herbivore resistance (reviewed in Lev-Yadun and Gould 2008; Naing and Kim 2021). For instance, reduced herbivory on morphs with higher anthocyanin pigmentation has been documented in *Raphanus sativus* (petal color variation, Irwin et al. 2003) and *Quercus coccifera* (leaf color variation, Karageorgou and Manetas 2006). Despite long-standing interest in the evolution of flower color very few studies have investigated the impact of stochastic process and gene flow on spatio-temporal variation in flower color (Schemske and Bierzychudek 2001, 2007; Turelli et al. 2001). Moreover, the possible interplay between neutral processes and selection on flower color variation across a gradient of urbanization remains poorly studied (Cabon et al. 2022).

Native to Eurasia, *H. matronalis* is an herbaceous biennial or short-lived perennial that was introduced to urban North America as an ornamental garden plant beginning in the 1800s and has since become naturalized across southern Canada and the northern United States (Rousseau 1968; Francis et al. 2009). The species is still widely sold across North America and naturalized populations are now particularly common in disturbed habitat along roads and railways, especially in urban areas (Francis et al. 2009). Populations of *H. matronalis* are often polymorphic for flower color, although the pattern of phenotypic variation in flower color has not been formally quantified. Majetic et al. (2007) recognized two color morphs, white and purple, and after a limited number of hand crosses reported that the pattern of color segregation was consistent with color being determined by a single Mendelian locus with white dominant to purple. However, others have noted that intermediate pink phenotypes are common within most populations (Mitchell and Ankeny 2001, Rothfels et al. 2002, Weeks & Frey 2007, Francis et al. 2009), which indicates that the inheritance of petal color in this species is likely more complex.

Little is known about the selective agents that may potentially act on flower color in *H. matronalis*. The species seems adapted to both day and night pollination (Francis et al. 2009) and flowers are visited by a variety of insects including bees, butterflies, moths and syrphid flies. Majetic et al. (2007, 2008) reported that insects pollinating *H. matronalis* do not exhibit flower color preferences and the floral scent composition, though variable and likely relevant to pollinators, is independent of color. The extent to which pollination is required for seed set is uncertain due to conflicting evidence that *H. matronalis* is self-compatible (Majetic et al. 2008) and possibly predominantly self-fertilizing (Susko and Clubb 2008) or self-incompatible and obligately outcrossing (Mitchell and Ankeny 2001; Weeks and Frey 2007). The composition of the herbivore fauna and whether any herbivore might respond to flower color is also unknown. Recently, an *H. matronalis* specialist weevil (*Ceutorhynchus inaffectatus* Gyllenhal, 1837, Curculionidae) from the native European range (Larsen et al., 1992) has been reported in southern Ontario (Pentinsaari et al., 2019) but it is not known whether it has any impact on introduced populations of *H. matronalis* or if it attacks plants non-randomly with respect to flower color.

We quantified pattern of flower color variation in *H. matronalis* and surveyed the frequency of three distinct flower color morphs in 136 stands across an urban to rural gradient in southeastern Ontario, Canada. To investigate the relative influence of neutral and selective processes on flower color variation in introduced populations of *H. matronalis* we tested whether flower color variation deviated from established theoretical expectations of a neutral cline. We tested five predictions consistent with a selectively neutral cline in flower color variation. (1) Flower color morph diversity should increase with stand size because larger stands are less vulnerable to the loss of genetic diversity via drift. Stands of this invasive species may be larger in urban than rural areas, which could contribute to a cline in increasing flower color diversity with increasing human activity. (2) Given that *H. matronalis* was and still is introduced via horticulture and that multiple introductions are most likely in urban areas, we predicted that flower color morph diversity would be greater in stands located in areas with higher levels of human activity and that this should be independent of the effect of stand size predicted above. (3) As a result of 1 and 2, variation in color morph diversity across generations should be lower in urban areas compared to rural areas, should increase with increasing temporal variation in stand size and should decrease with increasing mean stand size across generations. (4) Variation in color morph frequencies among stands across the urban-rural gradient should be random.

Reduced color diversity of rural stands should not be coincident with a systematic increase or decrease in the frequency of any one morph. Overall, the chance of a morph being stochastically lost from stands should be inversely proportional to its global frequency. (5) Color morphs should not differ in components or correlates of fitness.

Deviations from the neutral pattern of flower color variation described by these five predictions could indicate a possible role for selection. For example, if flower color morph diversity decreased with stand size (a departure from neutral prediction 1) this would be consistent with potentially more efficient selection in larger populations thereby reducing morph diversity via systematic change in morph frequencies. Any systematic change in morph frequencies may be reflected by fitness differences among morphs (a departure from neutral prediction 5). For example, fitness may be higher in more pigmented morphs, possibly due to higher anthocyanin levels deterring herbivores. If urban areas have a higher abundance of the introduced specialist weevil and therefore higher levels of herbivory, we predict that there may be a systematic increase in the frequency of the better-protected purple morph with urbanization (a departure from neutral prediction 4).

## METHODS

### Study area

We located 136 naturalized stands of *H. matronalis* over a 1300 km^2^ area within a 40 km radius of Kingston, Ontario, Canada in June 2021 and surveyed these stands over three consecutive years, which amounts to three generations for this biennial species (Appendix S1; see Supplemental Data with this article). The study area covered a gradient of human activity and contained stands that varied widely in census size. A stand was defined as a discrete group of plants separated by at least 100 m from other such groups (median distance was 500 m). We accessed and surveyed 132 of these stands in June 2021, 112 stands in June 2022 and 116 stands in June 2023. Of these, 105 stands were sampled in all three years. Five sites were devoid of plants after 2021; seven sites had plants in 2021 and 2023 but not 2022. For 19 stands, we could only gain access for one (12) or two (7) of the sampling years.

### Geographical survey of flower color variation

Previous work on flower color variation in *H. matronalis* has classified flowers as either light or dark (Majetic et al. 2007, 2008) though other researchers have noted considerable intermediate variation in flower color (Mitchell & Ankeny 2001, Rothfels et al. 2002, Weeks & Frey 2007). To better understand the pattern of phenotypic variation, we performed spectral analysis on more than 1000 flowers from three large stands of *H. matronalis* in our study area. This revealed enough intermediate flower color variation to justify a 3-morph classification as a more effective way of capturing (putatively) genetic variation in flower color within and among eastern Ontario stands (Appendix S2). For each stand, we estimated flower color morph frequencies by randomly sampling an average of 96 plants (range = 1–423) and scoring flowers as white (W), pink (P) or purple (V = violet) using a set of standard photographs to ensure consistency (Fig. 1). We sampled all plants in stands consisting of fewer than 100 plants and a random sample of 100 for larger stands and visually estimated the number of reproductive individuals (*N*) after surveying each site.

**Figure 1.**
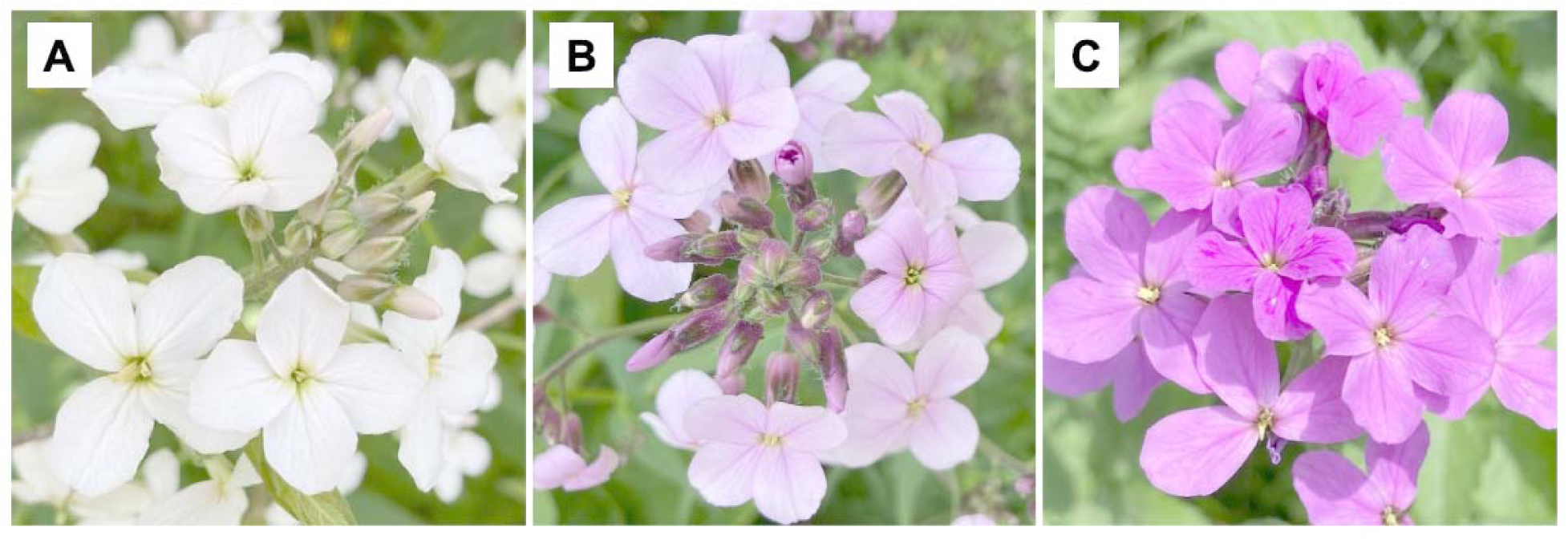
Flower color variation in introduced populations of *Hesperis matronalis* from eastern Ontario, Canada. White morph flowers (panel A) were pure white with no hints of pink. Pink flowers (panel B) ranged from having obvious hints of pink, often in the center of the flower to petals that were predominantly pink with regions of darker pink. Purple flowers (panel C) were solid purple.

We used night sky brightness as a proxy for the level of human activity around each sampled stand of *H. matronalis* (Sutton et al. 2001; Pun and So 2012). We obtained zenith sky brightness data from Falchi et al.’s (2016) world atlas of artificial sky luminance and accessed this data through lightpollutionmap.info. The zenith sky brightness for each stand was reported in units of magnitude per square arc second (mag arcsec^−2^) at a spatial resolution of 1 km.

Magnitudes are a measure of brightness and a smaller value indicates a brighter object. Zenith sky brightness ranges from 16 mag arcsec^−2^ for the lightest sky (high human activity) to 22 mag arcsec^−2^ for the darkest sky (low human activity). To make this measure easier to interpret in terms of human activity, we multiplied these values by –1 and refer to this as “night sky brightness” (NSB). As a result, there is a positive correlation between night sky brightness and human activity. For example, a rural stand could have an NSB value of –21, while an urban stand, where light pollution is greater, could have an NSB value of –19. Among the stands we sampled, NSB correlated strongly and negatively with distance from Kingston city center (Pearson *r* = – 0.82, *n* = 136, *P* < 0.0001).

### Comparing fitness correlates and seed predation among color morphs

In each of 22 populations surveyed in 2021, we tagged 10–20 randomly chosen plants per color morph. Then in mid-August, the above-ground portion of the 215 plants that could be relocated with intact tags were harvested when fruits were mature but seeds not yet released. After drying plants to constant mass at 70°C, we counted the number of mature fruits (each containing ≥ 1 filled seed) as well as the number of flowers that failed to develop fruits (indicated by persistent petioles). Above-ground plant size was estimated by total dry stem mass, because about 30% of plants had some of their leaves at this stage. We estimated flower number as the sum of fruit number and the number of fruits that did not develop. From each plant, we randomly selected five fruits and counted the number of filled seeds in each. Total seed production was estimated for each plant as the product of fruit number and the average number of filled, undamaged seeds per fruit. Many seeds were destroyed by the pre-dispersal seed predator *Ceutorhynchus inaffectatus* (D.J. Ensing, T. Nelson & C.G. Eckert, unpublished), so to estimate the number of seeds that could have potentially matured in the absence of seed predation, we counted the indentations left by developing seed in the silique septum. From these data, we estimated the proportion of seeds destroyed by *C. inaffectatus*. We also estimated potential seed production in the absence of seed predation as the product of fruit number and mean potential seeds/fruit.

### Statistical analyses

We measured flower color morph diversity of each stand using Shannon’s diversity index (*H*’) calculated using the *diversity* function in the vegan package (version 2.6-2; https://CRAN.R-project.org/package=vegan) for the R statistical environment (version 4.2.1, R Core Team 2022), which was used for all analyses. *H*’ varies from 0 (monomorphic) to 1.10 (all three morphs equally frequent) and is particularly appropriate when sample sizes are variable and sometimes small (Morris et al. 2014). *H*’ correlated strongly with other measures of morph diversity such as morph evenness (e.g. *r* ≥ +0.99, all *P* < 0.0001; Barrett et al. 1989). The distribution of stand size (*N*) was strongly right-skewed so we log_10_-transformed this variable for analysis (log_10_*N*). To test the predictions that flower color morph diversity would be higher in larger stands (higher log_10_*N*, prediction 1) and stands in areas with higher human activity (higher NSB, prediction 2), we fit variation in *H*’ to a linear model including log_10_*N*, NSB and their interaction as predictors using the *lm* function in base R. The response and predictor variables were scaled by their means and standard deviations to generate standardized partial regression coefficients that could be compared between predictors and years. We tested the significance of each predictor using backward stepwise elimination with likelihood ratio tests. Because a different number of sites were sampled in each year, we performed this analysis for each year separately. For all analyses, we ensured that residuals were normally distributed and independent from predicted values. As predicted, NSB and log_10_*N* correlated positively but weakly in all three years but the correlation was only significant for 2021 (2021: Pearson’s *r* = +0.35, *P* < 0.0001; 2022: *r* = +0.14, *P* = 0.13; 2023: *r* = +0.15, *P* = 0.11). We calculated the variance inflation factor (VIF) for both predictors using the *vif* function in the car package (version 3.1-1, https://CRAN.R-project.org/package=car) to confirm that collinearity between predictors did not complicate the interpretation of partial regression coefficients (VIF = 1.14 in 2021, 1.02 in 2022, 1.02 in 2023). We used added-variable plots to graphically display the direct effect of NSB and log_10_*N* on *H*’ while controlling the effect of the other predictor and accounting for the weak collinearity between predictors.

Variation in color morph diversity across generations (prediction 3) was evaluated using the 105 stands surveyed in all three years. To measure variation, we used the standard deviation of *H*’ across years instead of the coefficient of variation in *H*’ because the standard deviation was less correlated with mean *H*’ across years (standard deviation *r* = –0.31, coefficient of variation *r* = –0.85). We fit variation in the SD of *H*’ to a linear model with NSB, mean log_10_*N* and the SD of log_10_*N* as predictors, and evaluated the significance of predictors and displayed the data graphically as above. VIF’s for all predictors were < 1.1.

Stochastic processes involving genetic drift in small stands or random introduction of genotypes via horticulture should result in random variation in color morph frequencies across the urban-rural gradient. We analyzed two patterns in the data for departures from random variation (prediction 4). First, we tested whether there was a bias in which color morph predominated among stands that were fixed for a single morph (*H*’ = 0) and in which morph was lost from stands. Population genetic theory predicts that small populations will tend to fix (or lose) alleles more frequently than larger populations (Hedrick 2000), so we compared log_10_*N* between fixed and polymorphic populations using *t*-tests (*t.test* function in base R). In the specific case of *H. matronalis*, we also might expect urban populations to maintain all three color morphs more frequently than rural populations due to the higher frequency of multiple introductions in urban areas. Hence NSB should be higher in polymorphic than fixed populations. Theory also predicts that the probability of an allele being randomly fixed in a population is proportional to its global frequency (Hedrick 2000). Hence, we tested the neutral prediction that the frequency of *H. matronalis* stands stochastically fixed for a color morph should be proportional to the mean frequency of that morph in polymorphic populations using a chi-squared test with a *P* value derived from Monte Carlo simulation (*chisq.test* function in base R). Conversely, we tested whether the frequency of stands that had lost a particular morph was inversely proportional to the frequency of that morph in polymorphic populations.

Second, we investigated whether, contrary to neutral predictions, flower color morph frequencies varied systematically (i.e. nonrandomly) with human activity and stand size by fitting morph frequencies as a multinomial response variable to a generalized linear model using the *multinom* function in the nnet R package (version 7.3-17, https://CRAN.R-project.org/package=nnet). We included NSB and log_10_*N* and their interaction as fixed predictors and evaluated significance with stepwise likelihood ratio tests as above. Because the interaction was significant for both years, we illustrated trends graphically by plotting model predictions made using the *predict* function in base R over the full range of NSB and at the 25-percentile, median and 75-percentile of log_10_*N*. We determined how well the model predicted morph frequencies for each morph separately by regressing observed frequencies over predicted frequencies and calculating the *r*^2^ value and *P* value from an *F*-test.

We found a systematic change in color morph frequencies with both NSB and log_10_*N* involving an increase in the frequency of the purple morph (and a corresponding decrease in frequency of the pink and, to a lesser extent, the white morphs) in stands that were smaller and in areas of lower human activity. The increase in the frequency of the purple morph correlated negatively with morph diversity in all years (2021 *r* = –0.63, 2022 *r* = –0.68, 2023 *r* = –0.67, all *P* < 0.0001). This result led us to test whether human activity and stand size influenced morph diversity even after accounting for the systematic increase in the frequency of the purple morph, that is whether stochastic processes might affect morph diversity above and beyond the systematic increase in the frequency of the purple morph. To do this, we used the morph frequency values predicted by our multinomial regression to calculate ‘expected morph diversity’ (*H’*_pred_) and included this as a covariate in the previously described linear models. All together this allowed us to examine the effect of human activity (NSB) and stand size (log_10_*N*) on observed morph diversity (*H*’) while accounting for the impact of individual morph frequency on diversity. We performed significance tests, checked assumptions and made graphical displays of the results as described above for the original regression analysis. There was some collinearity among predictors as expected because *H’*_pred_ correlated positively with both NSB (2021 *r* = +0.68, 2022 *r* = +0.63, 2023 *r* = +0.38, all *P* < 0.0001) and log_10_*N* (2021 *r* = +0.77, 2022 *r* = +0.65, 2023 *r* = +0.39, all *P* < 0.0001), however the VIF was < 5 for all predictors in all years (2021: NSB = 2.12, log_10_*N* = 2.80, *H’*_pred_ = 4.53; 2022: NSB = 2.10, log_10_*N* = 2.16, *H’*_pred_ = 3.55; 2023: NSB = 1.17, log_10_*N* = 1.18, *H’*_pred_ = 1.36).

To compare fitness components and seed predator damage among color morphs (prediction 5), we fit variation in several fitness related response variables to linear models with color morph, NSB and their interaction as fixed effects and stand as a random effect and evaluated significance with stepwise likelihood ratio tests as above. We fit log_10_-transformed above-ground stem dry mass to a linear mixed-effects model using the *lmer* function in the lme4 R package (version 1.1-30, https://CRAN.R-project.org/package=lme4). Because flower number, fruit number, seed number, and potential seed number per plant are over-dispersed count data we used generalized linear mixed-effects models (GLMM) with negative binomial errors fit using the *glmmTMB* function in the glmmTMB package (version 1.1.4, https://cran.r-project.org/package=glmmTMB). The negative binomial distribution is appropriate for count data and, unlike the Poisson distribution, includes a dispersion parameter. For the proportion of flowers setting fruit and the proportion of seeds destroyed by *Ceutorhynchus inaffectatus* we used GLMM with binomial errors fit with *glmmTMB*. We evaluated whether the distribution of residuals fit model assumptions using the *simulateResiduals* function and tested for over/under dispersion of residuals using the *testDispersion* function, both in the DHARMa package (version 0.4.6, https://cran.r-project.org/package=DHARMa). Residuals for fruit set were over dispersed, so we modeled a dispersion parameter using the *dispformula* argument in *glmmTMB*. Seeds per plant was modeled with a zero-inflation parameter applied to all observations using the *ziformula* argument.

## RESULTS

### Variation in color morph diversity

There was wide variation in the frequencies of the three flower color morphs among stands in all years (Fig. 2). Flower color morph diversity (*H*’) varied across almost the full range of possible values in 2021 (range = 0–1.09, mean = 0.71), 2022 (range = 0–1.10, mean = 0.77) and 2023 (range = 0–1.10, mean 0.81). As predicted, flower color morph diversity increased significantly with both stand size (log_10_*N*, prediction 1) and human activity (NSB, prediction 2) in all years (Table 1, Fig. 3) and the effects of these two predictors did not interact (all *P* > 0.21).

**Figure 2.**
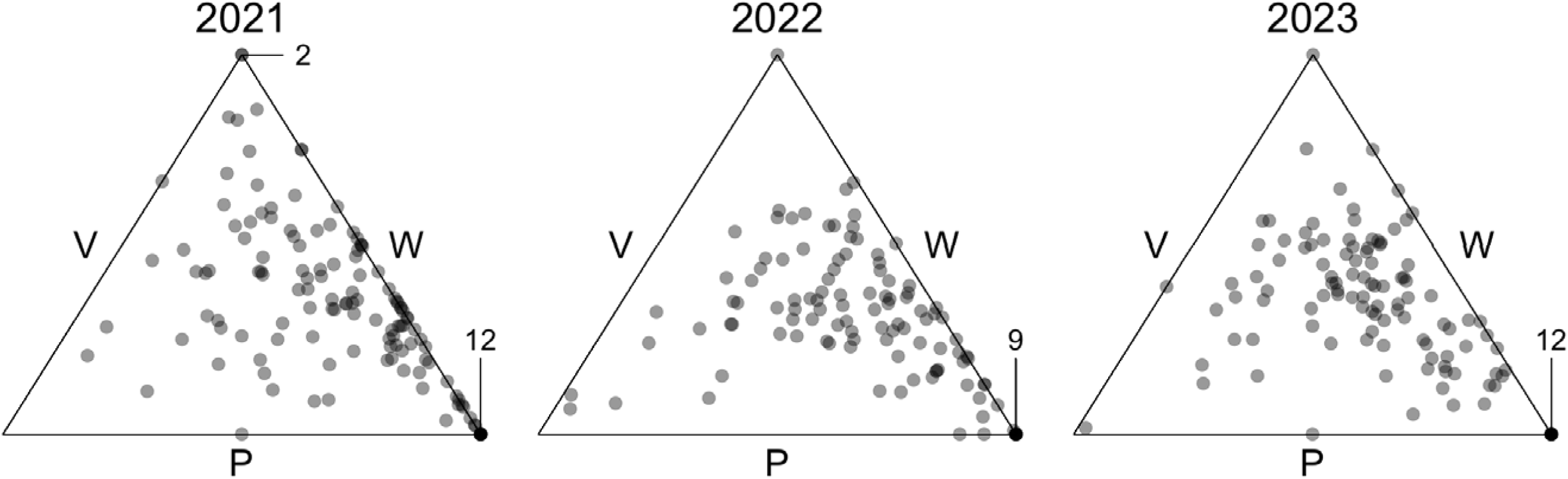
Variation in color morph frequencies among stands of *Hesperis matronalis* in eastern Ontario, Canada. Each point represents the morph frequencies in a stand (132 stands in 2021, 112 in 2022, 116 in 2023). Color morphs are white (W), pink (P) and purple (V and the distance of a point from a morph’s side is proportional to the frequency of that morph in the stand. Points in the center of the triangle have equal frequencies of all three morphs, those lying on a side lack that morph, and those lying on a vertex are fixed for the morph indicated on the opposite side. Lines and numbers indicate the number of points overlapping on a vertex.

**Figure 3.**
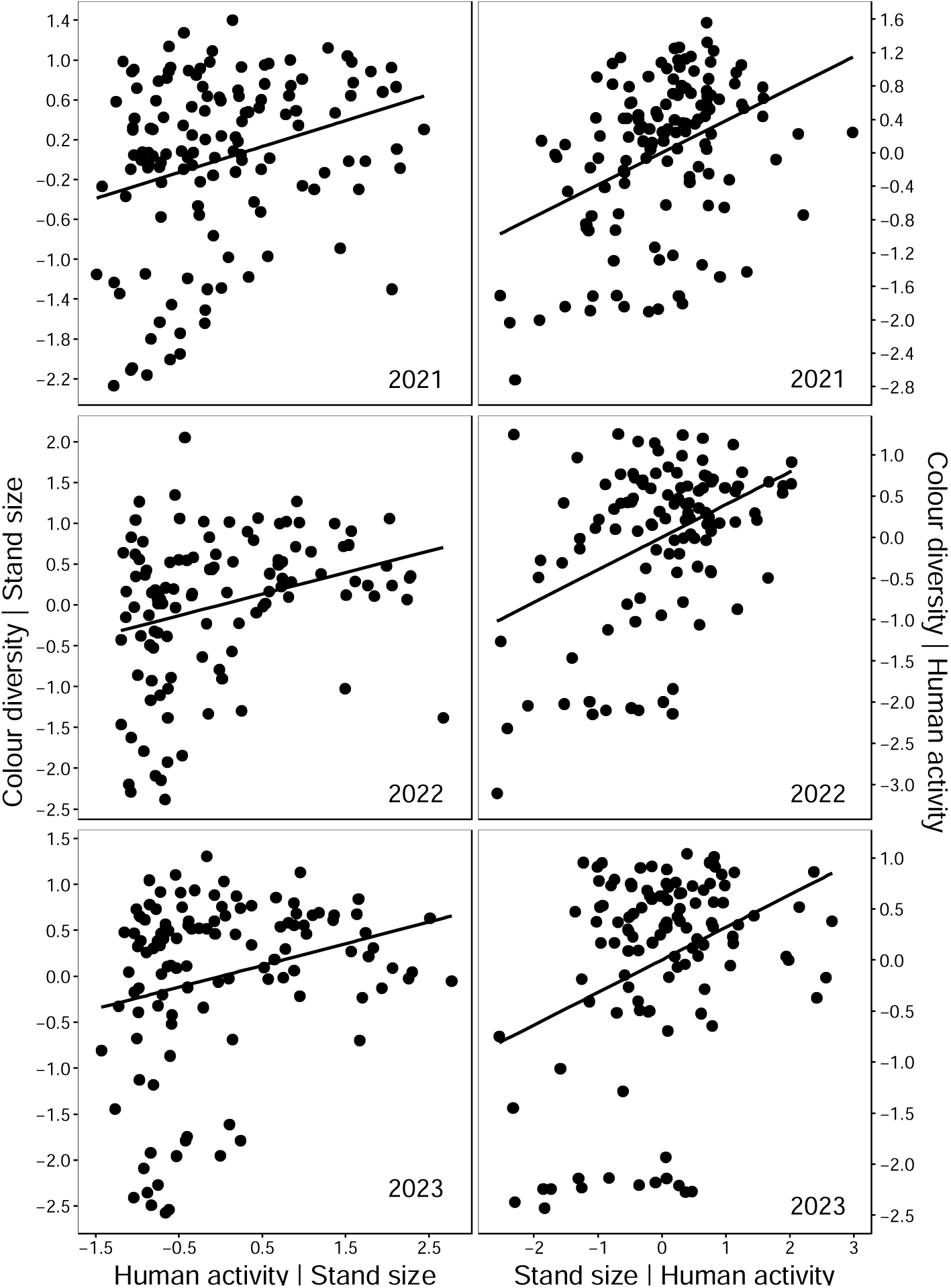
Added-variable plots showing the relations between flower color morph diversity (*H’*) and both human activity (NSB) and stand size (log_10_*N*) among stands of *Hesperis matronalis* from eastern Ontario, Canada sampled over three years (132 stands in 2021, 112 in 2022, 116 in 2023). Each point is a stand. The y-axis is residual *H’* when the other predictor is held constant. The x-axis is the residual of the focal predictor when the other predictor is held constant.

**Table 1.** Analyses of variation in color morph diversity (*H*’) among stands of *Hesperis matronalis*. In analysis A, variation in *H’* was fit to linear models with human activity (NSB), stand size (log_10_*N*) and their interaction as predictors (132 stands for 2021, 112 for 2022, 116 for 2023). Cells include *F*-tests of significance and standardized partial regression coefficients (*b*) when significant, or NA when that predictor was not included in that analysis. The effect of both human activity and stand size on *H’* (Fig. 2) might have been due to a systematic increase in the frequency of the purple morph in smaller stands and in areas with low human activity. In analysis B we tried to control for this effect by entering “predicted morph diversity” (*H’*_pred_) calculated from predicted morph frequencies (Fig. 3) as a covariate in the linear models. *H’*_pred_ was only a significant predictor of *H’* in 2023 and this reduced the effect of both NSB and log_10_*N*. However, predicted morph diversity correlated with both these other predictors, resulting in relatively high variance inflation factors (see Results).

|  | A |  |  | B |  |  |
| --- | --- | --- | --- | --- | --- | --- |
| Year = | 2021 | 2022 | 2023 | 2021 | 2022 | 2023 |
| $r^2$ = | 0.29 | 0.26 | 0.18 | 0.29 | 0.26 | 0.21 |
| Predictor |  |  |  |  |  |  |
| Human activity<br>(NSB) | $b = +0.261$<br>$F_{1,129} = 10.7$<br>$P = 0.0013$ | $b = +0.264$<br>$F_{1,109} = 10.0$<br>$P = 0.0020$ | $b = +0.236$<br>$F_{1,113} = 7.5$<br>$P = 0.0071$ | $b = +0.261$<br>$F_{1,129} = 10.7$<br>$P = 0.0013$ | $b = +0.264$<br>$F_{1,109} = 10.0$<br>$P = 0.0020$ | $b = +0.170$<br>$F_{1,112} = 3.5$<br>$P = 0.064$ |
| Stand size ($\log_{10}N$) | $b = +0.384$<br>$F_{1,129} = 23.3$<br>$P < 0.0001$ | $b = +0.396$<br>$F_{1,109} = 22.5$<br>$P < 0.0001$ | $b = +0.319$<br>$F_{1,113} = 13.7$<br>$P = 0.00033$ | $b = +0.384$<br>$F_{1,129} = 23.3$<br>$P < 0.0001$ | $b = +0.396$<br>$F_{1,109} = 22.5$<br>$P < 0.0001$ | $b = +0.251$<br>$F_{1,112} = 7.5$<br>$P = 0.0070$ |
| NSB x $\log_{10}N$ | —<br>$F_{1,128} = 0.30$<br>$P = 0.58$ | —<br>$F_{1,108} = 1.56$<br>$P = 0.21$ | —<br>$F_{1,112} = 0.20$<br>$P = 0.66$ | NA | NA | NA |
| Predicted morph<br>diversity ($H'_{\text{pred}}$) | NA | NA | NA | —<br>$F_{1,128} = 1.9$<br>$P = 0.17$ | —<br>$F_{1,108} = 0.42$<br>$P = 0.52$ | $b = +0.199$<br>$F_{1,112} = 4.15$<br>$P = 0.044$ |

Among the 105 stands sampled in all three years, there was year to year consistency in both stand size and color morph diversity, as *H*’ corelated strongly between years (Kendall’s coefficient of concordance (*W*) = +0.75, *P* < 0.0001) and so did log_10_N (*W* = +0.80, *P* < 0.0001). The correlation between *H*’ and both NSB and log_10_*N* varied slightly between years but was always positive (Appendix S3). On average, *H*’ varied among years and was significantly higher in 2023 than in the two previous years (mean ± 1SE: 2021= 0.734 ± 0.031, 2022 = 0.768 ± 0.032, 2023 = 0.847 ± 0.029). This was associated with significantly higher stand size in 2023. For individual stands, the increase in flower color morph diversity between 2023 and the two previous years was higher for stands in areas with lower human activity (lower NSB). This increase correlated positively but not significantly with the between-year increase in stand size and was not related to the year-to-year change in sample size (Appendix S4).

Consistent with prediction 3, the among-generation variation in *H*’ (Appendix S5) was lower in stands that were: a) larger (mean of log_10_*N* across generations: standardized partial regression *b* –0.31, *F*_1,101_ = 11.3, *P* = 0.0011), b) less variable in size (SD of log_10_*N* across generations: *b* = +0.29, *F*_1,101_ = 10.1, *P* = 0.0019), and c) in areas with higher human activity, though this effect was not significant (NSB: *b* = –0.13, *F*_1,101_ = 2.0, *P* = 0.16).

### Nonrandom variation in morph frequencies

The stands we sampled were usually polymorphic for flower color (89.4% of 132 in 2021; 91.1% of 112 in 2022; 88.8% of 116 in 2023), and most of these polymorphic stands included all three color morphs (73.7% in 2023; 88.2% in 2022; 93.2% in 2023). Consistent with neutral expectations, monomorphic stands were about a tenth of the size of polymorphic stands and tended to be in areas with lower human activity (Table 3). In contrast to neutral expectations, monomorphic stands were almost always fixed for the purple morph. Fixed purple stands made up 86% of 14 monomorphic stands in 2021, 90% of 10 in 2022 and 92% of 13 in 2023. This is significantly higher than would be expected based on the mean frequency of the purple morph in polymorphic populations (0.501 in 2021 χ^2^ = 7.10, simulated *P* = 0.019; 0.495 in 2022 χ^2^ = 6.56, *P* = 0.011; 0.428 in 2023 χ^2^ = 13.01, *P* = 0.00050). Moreover, all five stands that were monomorphic in all three years were fixed for purple (Table 2).

**Table 2.** Variation in frequency of color morphs in stands of *Hersperis matronalis* from eastern Ontario, Canada surveyed in three consecutive years (generations). Average morph frequency was calculated across all stands and from only stands polymorphic for flower color. The bottom half of the table shows the numbers of populations fixed for each color morph in each year and all years.

| Subset | Year | n stands | Color morph frequency |  |  |
| --- | --- | --- | --- | --- | --- |
|  |  |  | White | Pink | Purple |
| All stands | 2021 | 132 | 0.129 | 0.333 | 0.538 |
|  | 2022 | 112 | 0.174 | 0.295 | 0.531 |
|  | 2023 | 116 | 0.192 | 0.324 | 0.484 |
| Polymorphic stands | 2021 | 118 | 0.144 | 0.355 | 0.501 |
|  | 2022 | 102 | 0.190 | 0.314 | 0.495 |
|  | 2023 | 103 | 0.216 | 0.356 | 0.428 |
|  |  |  | Number of stands fixed for: |  |  |
|  |  |  | White | Pink | Purple |
| Monomorphic stands | 2021 | 14<sup>1</sup> | 0 | 2 | 12 |
|  | 2022 | 10<sup>2</sup> | 0 | 1 | 9 |
|  | 2023 | 13<sup>3</sup> | 0 | 1 | 12 |
|  | All 3 years | 5<sup>4</sup> | 0 | 0 | 5 |

Notes:
(1) Five of these stands were gone in 2022, two of which reappeared in 2023. Of the 11 resampled stands, four were polymorphic in future years, one just barely. Five stands had < 5 plants but five had > 30 plants.
(2) Six of these stands were also fixed in 2021 (five of which stayed fixed in 2023). Three were polymorphic in 2021. One was located in 2021 but only became accessible in 2022 and remained fixed in 2023. Three stands had < 5 plants but four had > 30 plants.
(3) Nine of these stands were fixed in a previous year (five for all three years). Four were polymorphic in a previous year (including two that were gone in 2022). Two stands had < 5 plants but six had > 30 plants.
(4) The mean size of these stands across all three years ranged 19.3 – 144.0 and averaged 74.1.

**Table 3.** Comparison of stand size (log_10_*N*) and human activity (NSB) between populations of *Hesperis matronalis* monomorphic vs. polymorphic for flower color. Monomorphic stands made up 14 of 132 stands in 2021, 10 of 112 stands in 2022 and 13 of 116 stands in 2023. Monomorphic and polymorphic stands were compared using Welch’s 2-sample *t*-test with a 1-tailed test of significance.

| Variable | Year | Monomorphic | Polymorphic | $t$-test (1-tailed $P$) |
| --- | --- | --- | --- | --- |
| Stand size<br>$\log_{10}N$ | 2021 | $1.19 \pm 0.17$ | $2.11 \pm 0.06$ | $t = 5.08, P < 0.0001$ |
| | 2022 | $1.29 \pm 0.18$ | $2.10 \pm 0.06$ | $t = 4.21, P = 0.0007$ |
| | 2023 | $1.45 \pm 0.21$ | $2.26 \pm 0.07$ | $t = 3.63, P = 0.0012$ |
| Human activity<br>NSB | 2021 | $-21.5 \pm 0.14$ | $-20.9 \pm 0.07$ | $t = 3.73, P = 0.0006$ |
| | 2022 | $-21.4 \pm 0.26$ | $-21.0 \pm 0.08$ | $t = 1.50, P = 0.080$ |
| | 2023 | $-21.6 \pm 0.07$ | $-21.0 \pm 0.08$ | $t = 6.10, P < 0.0001$ |

When stands lacked one morph (23.5% of stands in 2021, 10.7% in 2022, 6.0% in 2023), they usually lacked the white morph (Fig. 2; 93.5% of 31 dimorphic stands in 2021, 83.3% of 12 in 2022, 71.4% of 7 in 2023). However, this did not differ from neutral expectations based on the relative low frequency of the white morph in polymorphic populations (Table 2; 0.144 in 2021 χ^2^ = 1.59, simulated *P* = 0.29; 0.190 in 2022 χ^2^ = 0.04, *P* > 0.9; 0.216 in 2023 χ^2^ = 0.20, *P* > 0.9).

In contrast to neutral expectations, multinomial logistic regression revealed that the frequencies of the three color morphs covaried with both human activity (NSB) and stand size (log_10_*N*) and these effects interacted (Fig. 4; Table 4). In particular, the purple morph decreased in frequency with increasing human activity in all years. This was associated with opposing but somewhat weaker and less consistent increases in the frequencies of the pink and white morphs. The effect of stand size (log_10_*N*) on morph frequencies was less consistent and depended on the level of human activity.

**Figure 4.**
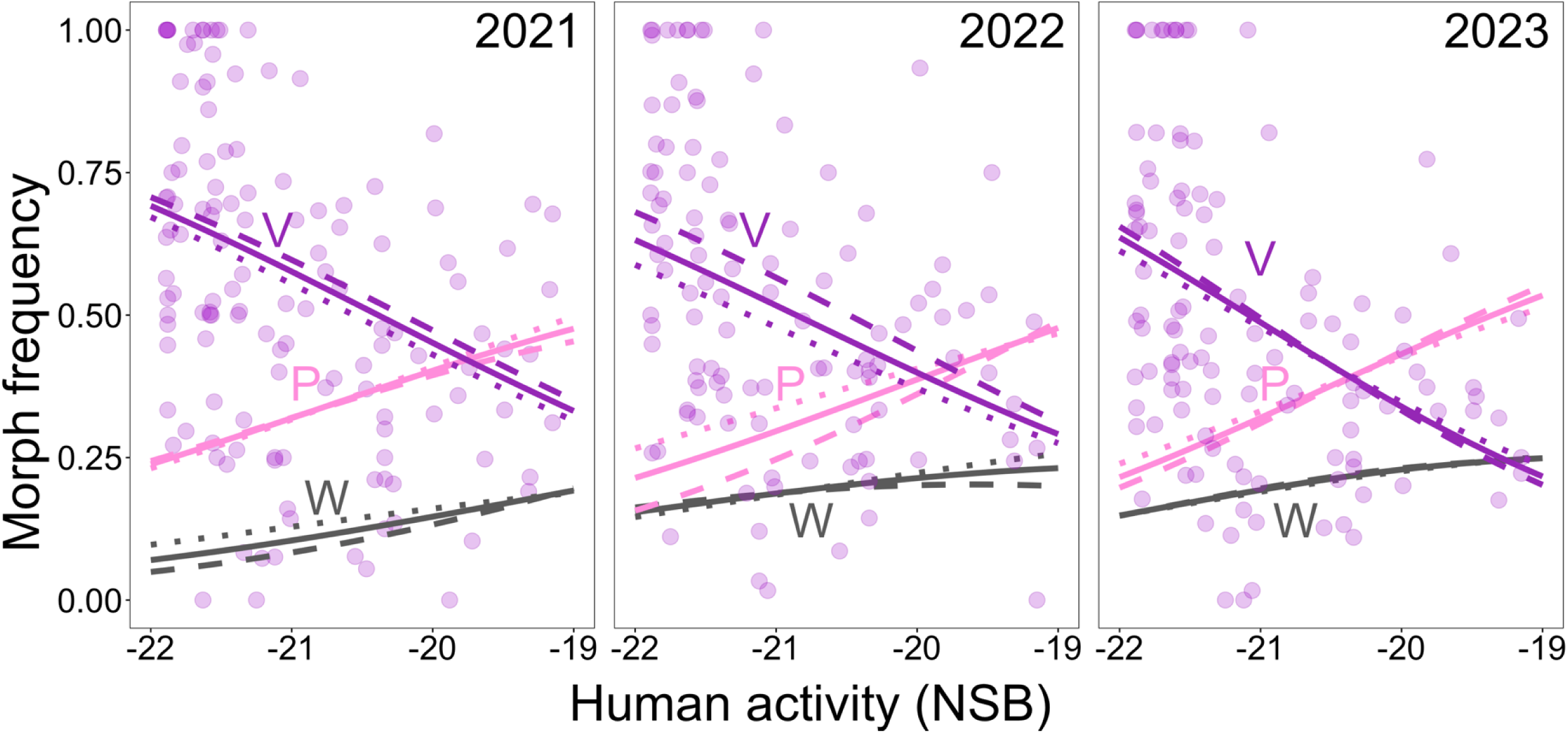
Systematic changes in flower color morph frequencies with both human activity (NSB) and stand size (log_10_*N*) among stands of *Hesperis matronalis* in eastern Ontario, Canada sampled over three years (132 stands in 2021, 112 in 2022, 116 in 2023). NSB and log_10_*N* interacted in their effects on morph frequencies (Table 3). Lines show the multinomial regression predictions for each color morph (W = white, P = light pink, V = purple). Solid lines show predictions at the median log_10_*N*, dashed lines at the 25^th^ percentile and dotted lines at the 75^th^ percentile. Although the effect of log_10_*N* on the frequency of each morph varies with NSB, the effect of NSB is more consistent. The points show frequencies of the purple morph to emphasize the relatively weak predictive ability of these regressions (*r*^2^ values in Table 3).

**Table 4.** Analysis of variation in flower color morph frequencies among stands of *Hesperis matronalis* in eastern Ontario, Canada. The numbers of the three color morphs sampled in each stand were modeled as a multinomial response variable with human activity (NSB) and stand size (log_10_*N*) and their interaction as predictors. For all three years, the best model included the interaction. The bottom half of the table reports the strength of regressions of observed morph frequencies on morph frequencies predicted by the model.

| Year | 2021 |  | 2022 |  | 2023 |  |
| --- | --- | --- | --- | --- | --- | --- |
| Multinomial regression |  |  |  |  |  |  |
| Predictor | LRT $\chi^2$ | $P$ | LRT $\chi^2$ | $P$ | LRT $\chi^2$ | $P$ |
| Human activity (NSB) | 373.3 | < 0.0001 | 374.1 | < 0.0001 | 567.2 | < 0.0001 |
| Stand size ($\log_{10}N$) | 73.5 | < 0.0001 | 154.8 | < 0.0001 | 10.5 | 0.0053 |
| NSB x $\log_{10}N$ | 30.5 | < 0.0001 | 31.0 | < 0.0001 | 19.6 | < 0.0001 |
| Observed vs. predicted morph frequencies |  |  |  |  |  |  |
| | $r^2$ | $P$ | $r^2$ | $P$ | $r^2$ | $P$ |
| White | 0.134 | < 0.0001 | 0.041 | 0.033 | 0.039 | 0.034 |
| Pink | 0.082 | 0.00083 | 0.271 | < 0.0001 | 0.169 | < 0.0001 |
| Purple | 0.177 | < 0.0001 | 0.230 | < 0.0001 | 0.173 | < 0.0001 |

Including color morph diversity predicted from this systematic variation in morph frequencies (*H’*_pred_) as a covariate in the linear models examining the effect of NSB and log_10_*N* on *H*’ weakened the effect of both these predictors only in 2023 (Table 1). *H’*_pred_ was not a significant predictor of *H’* in 2021 or 2022.

### Variation in seed predation but not size or seed production among morphs

The plants we sampled in 2021 varied dramatically in size and reproductive success. Dry stem mass varied 100-fold among individuals (0.68–72.65g) with a median of 5.57g. Flowers produced per individual varied more than 50-fold (11–736 flowers, median = 67). The number of filled seeds per plant ranged from 0–10,146 (median = 378). Despite this wide variation, none of these measures differed among color morphs (Table 5). The proportion of flowers forming fruit (fruit set) declined slightly with increasing human activity for the pink and purple morphs but not the white morph (Fig. 5). However, fruit set was generally high (mean = 0.95, fruit set = 1 for 48% of plants) and did not vary among morphs on average. Much of the variation in seed production among plants was due to pre-dispersal seed predation by *Ceutorhynchus inaffectatus*. Predation was near ubiquitous as all plants had at least one of five sampled fruits damaged, and 73.0% of plants suffered damage to all five fruits. All individuals had some seed destroyed, and the proportion of seeds destroyed varied from 0.36–1.00 (median = 0.76). Although this seed predator was also introduced from Europe and thus could be potentially more prevalent in areas with higher human activity, the proportion of seeds destroyed generally decreased with increasing human activity. The decrease was steeper for the purple and pink morphs than the white morph, and contrary to predictions, mean seed predation was highest in the purple morph (∼6% higher than among white morph plants) and intermediate in the pink morph. Although, variation in seed predation among morphs was not reflected in differences in per-capita realized seed production among morphs, estimated potential seed production per plant in the absence of predation increased by 135% with increasing urbanization for the white morph and decreased by 36% for the pink and purple morphs. As a result, the pink and purple morphs had a higher potential for seed production than the white morph in rural habitat and lower in urban habitat (Fig. 5).

**Figure 5.**
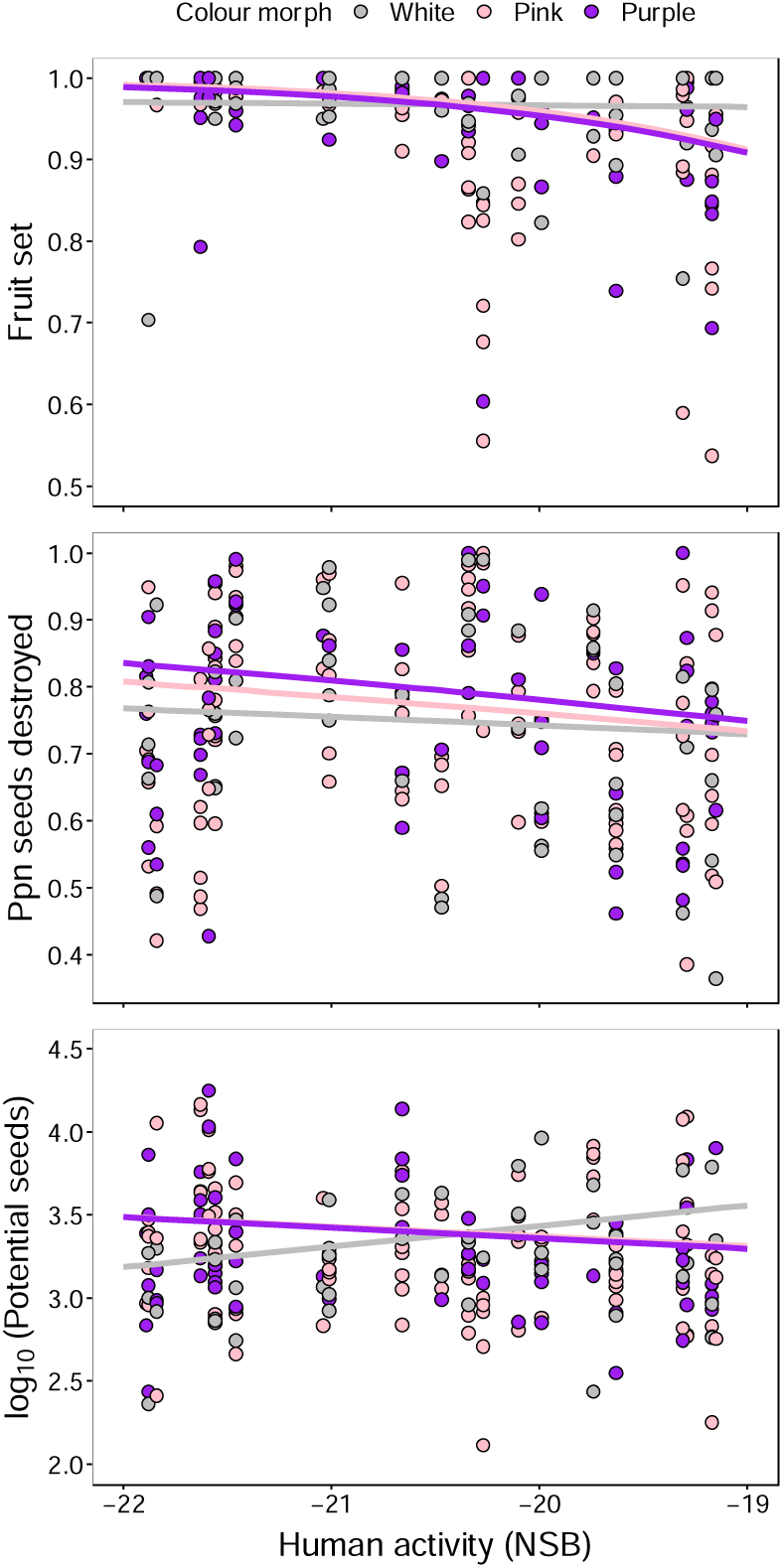
Variation in the proportion of flowers forming fruit (fruit set) and the proportion of seeds destroyed by the weevil seed predator *Ceutorhynchus inaffectatus* and the estimated potential seeds per plant in the absence of predation across a gradient of human activity and among flower color morphs in stands of *Hesperis matronalis* from eastern Ontario, Canada. Points are individual plants. Sample sizes were: white = 47 plants (grey points), pink = 109 (pink points), purple = 59 (purple points). Analysis of these data is in Table 5. Potential seeds per plant is log_10_-transformed for display only. The regression of fruit set and potential seeds over NSB varied significantly among morphs but mean fruit set or potential seeds did not vary among morphs. The regression of proportion seeds destroyed over NSB also varied among morphs and was, on average, higher for the purple morph, lower for the pink morph and lower still for the white morph. No other fitness measures varied with NSB or among morphs.

**Table 5.**
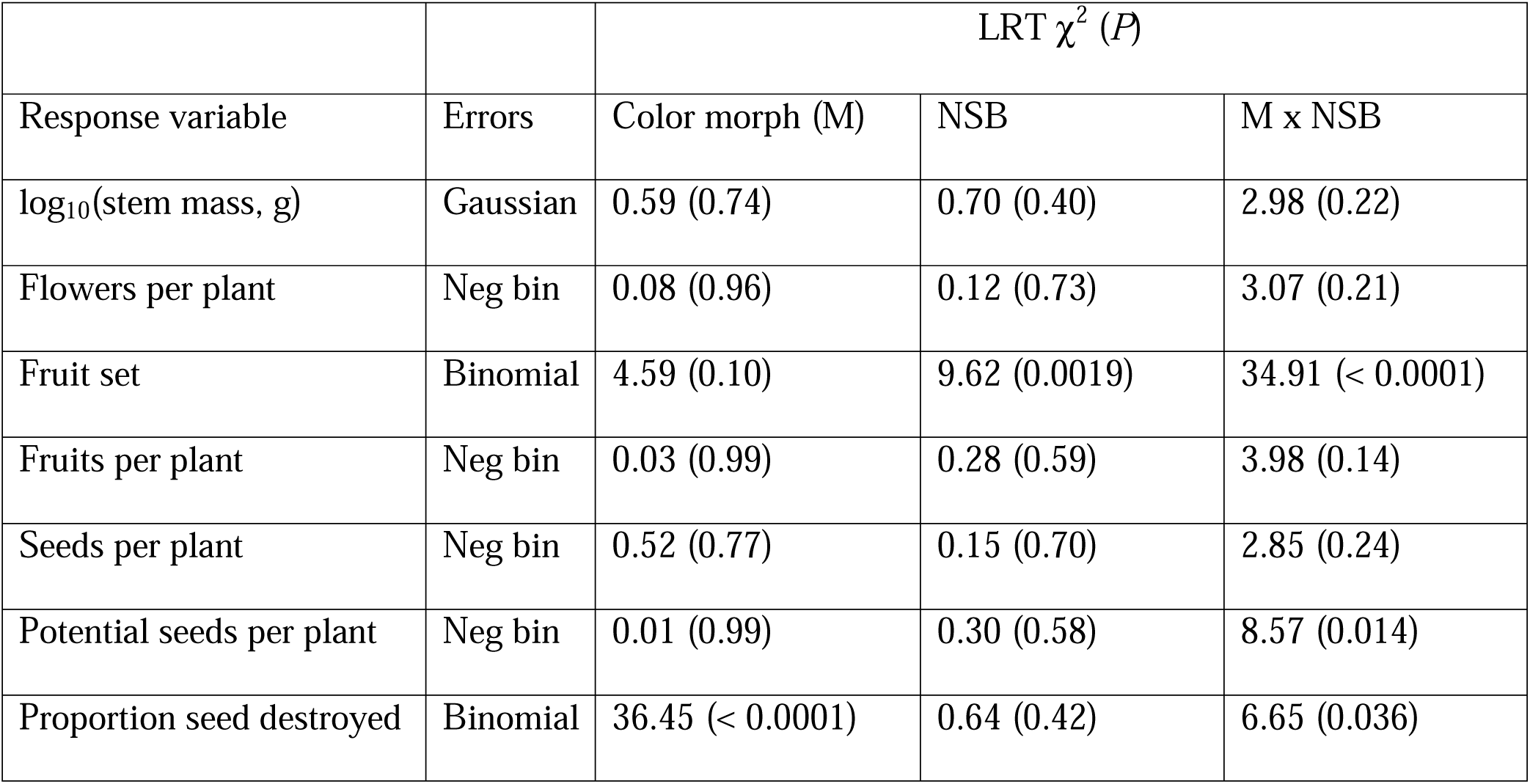
Analysis of variation in plant size, flower, fruit and seed production and seed predation among flower color morphs of *Hersperis matronalis* across 22 sites along a gradient of human activity (NSB) in eastern Ontario, Canada. Variation in each response variable was fit to a generalized linear model with color morph (df = 2), NSB (df = 1) and their interaction (df = 2) as fixed effects, site as a random effect, and an error distribution appropriate for the response variable (Errors, Neg bin = negative binomial). We evaluated significance of color morph using chi-squared likelihood ratio tests (LRT). Data for fruit set (proportion of flowers forming fruit) and proportion seeds destroyed are displayed in Fig. 5.

## DISSCUSSION

We investigated the evolutionary forces influencing flower color variation among 136 stands of an introduced biennial plant along an urban-rural gradient in eastern Ontario, Canada over three consecutive generations. Taken together, our results are consistent with color morph frequencies being influenced by the interplay of genetic drift and gene flow. In line with neutral predictions 1 and 2, color morph diversity was higher in larger stands and in areas of higher human activity where horticultural introductions and the dispersal of seeds are likely more frequent. Moreover, among-generation variation in color diversity was lower in urban areas compared to rural areas and in large stands compared to small stands (neutral prediction 3). However, in contrast to neutral prediction 4, we detected non-random variation in morph frequencies. The purple morph was more common in smaller stands and in rural areas, and this was associated with the fixation of the purple morph in some rural stands. Moreover, these fixed stands were not geographically clustered, suggesting independent fixation events (Appendix S6). Yet, the morphs did not differ in above-ground size, flower production, or lifetime seed production (neutral prediction 5).

Unexpectedly, purple morph plants suffered the most damage by seed predators and this damage decreased with increasing human activity. However, variation in seed predation was not reflected in differences among morphs in total seed production. Accounting for the non-random trends in morph frequencies only slightly (and usually non-significantly) reduced the associations between color morph diversity and both stand size and human activity.

In agreement with neutral expectations, flower color morph diversity (*H*’) correlated positively with stand size in all three generations. Similarly, the between-generation variation in *H*’ was lower in stands that maintained a larger mean size or exhibited less variation in size across generations. Because *H*’ likely represents genetic variation at the loci responsible for flower color, this is consistent with the expectation that genetic diversity is higher in larger populations, which has been supported by analyses of putatively neutral protein and DNA sequence variation in diverse species, especially plants (Frankham 1996; Leimu and Fischer 2008). Population size also affects genetic variation underlying morphological polymorphisms that may potentially influence fitness, for instance style morph frequencies in heterostylous plants where morph frequencies are under negative frequency-dependent selection (Eckert et al. 1996; Costa et al. 2016). Similarly, population size also correlated positively with morphological variation in *Salvia pratensis* and *Scabiosa columbaria* (Ouborg et al. 1991). Although analysis of variation in flower color morph frequencies, specifically in *Linanthus parryae*, was involved in some of the first tests of neutral processes in nature (Epling and Dobzhansky 1942; Schemske and Bierzychudek 2001), few studies have since investigated the relation between population size and flower color morph diversity across space or time (Sapir et al. 2021). In *Cirsium palustre*, morph frequencies did not systematically vary with population size, but increased population size from year to year was associated with increased flower color morph diversity (Mogford 1972, 1974). A small number of studies have looked at the relation between population size and the composition of color morph frequencies within populations, without considering morph diversity (Jersáková et al. 2006; Imbert 2021).

Habitat fragmentation associated with urbanization often reduces population size and gene flow, thereby making urban populations more vulnerable to the loss of genetic variation via genetic drift (Bartlewicz et al. 2015; Munshi-South et al. 2016; Miles et al. 2019). In contrast, we predicted larger population size and higher color morph diversity in urban areas because the introduction and spread of non-native species, particularly plants used in horticulture, are associated with human activity (Barney 2006; Pairon et al. 2010). Among the stands of *H. matronalis* we sampled, log-transformed stand size correlated positively, though not strongly with human activity (measured as NSB, *r* ≤ +0.35, see methods), and human activity positively influenced color morph diversity independently of variation in stand size. *Hesperis matronalis* was introduced into North America as a garden ornamental more than 200 years ago and continues to be sold for that purpose (Francis et al. 2009). It is likely that there have been multiple independent introductions of *H. matronalis* from gardens into naturalized populations. We searched the internet and found 26 North American seed retailers that currently sell *H. matronalis*, and 46% of these advertised their seed stock as including more than one flower color (Appendix S7). It is possible that urban gardens are planted with genetically diverse seed stock and may serve as a source of flower color diversity for naturalized populations.

An alternative explanation, not mutually exclusive of multiple introductions, is that urban populations of this invasive species are in closer proximity to one another than rural populations, which would enhance the maintenance of genetic variation at flower color loci via gene flow. Field observations made during three years of sampling *H. matronalis* in eastern Ontario clearly indicated that stands are much more frequent and closer together in urban than rural areas, as has been observed with other introduced species (Johnson et al. 2018). However, quantitative evidence of this in *H. matronalis* is lacking. Records of the species on iNaturalist (inaturalist.ca) are clearly more frequent closer to Kingston city center (especially if the records from our systematic survey are excluded), but this is likely due to a general trend of more iNaturalist posts and contributors closer to urban areas (Di Cecco et al. 2021).

Contrary to neutral expectations, we observed that morph frequencies varied systematically with human activity. The purple morph increased in frequency in areas with lower human activity and was fixed in more rural populations than would be expected by chance. If flower color was a neutral trait, theory would predict that the probability of any color morph becoming fixed is equal to the frequency of each morph in the global population (Hedrick 2000). In contrast, more than 80% of fixed populations were fixed for the purple morph despite the frequency of the purple morph being 40–50% depending on the study year. It is possible that the observed spatial pattern in morph frequency is simply the result of historical contingency (Eckert and Barrett 1995; Koski and Galloway 2020). Of the retailers that sell *H. matronalis* seeds, 50% appear to offer seed stock that is monomorphic for flower color (Appendix S7). It is possible that there was an early introduction of purple morph seeds to the rural end of our urbanization gradient. However, if the dominance of the purple morph in rural areas is due to happenstance we might expect to see spatial clustering of populations with a high proportion of purple flowers around the site of the historical accident. However, stands that were fixed for or have a high frequency of the purple morph were not obviously clustered (Appendix S6). Therefore, historical contingency is not a compelling explanation for non-random variation in morph frequency.

Most studies of flower color clines have focussed on geographic variation in selective pressures, particularly pollinator interactions (Sapir et al. 2021). Of course, other biotic and abiotic factors, such as herbivores and climate, can also exert selection on patterns of flower color variation (Strauss and Whittall 2007; Caruso et al. 2018; Koski and Galloway 2020).

However, our analysis of variation in plant size, seed predation and seed production among morphs, albeit with modest and unbalanced samples, suggests that the dominance of the purple morph in rural populations of *H. matronalis* is not easily explained by differential selection of the color morphs along the urban-rural gradient. Most measures of plant performance did not vary among color morphs or with human activity. We did find that the fruit set of purple and pink morphs, but not white, decreased with increasing human activity and that the color morphs differed in pre-dispersal seed predation, particularly at rural sites. However, these effects were not reflected in differences among morphs in lifetime seed production (see also Mitchell and Ankeny 2001; Weeks and Frey 2007).

An important caveat for our comparison of fitness correlates among morphs is that we have only measured lifetime seed fitness. Thus, it remains possible that there is a significant difference between morphs in pollen fitness through either difference in pollen production or pollinator visitation, which is especially relevant because *H. matronalis* is likely predominantly outcrossing (Mitchell and Ankeny 2001; Weeks and Frey 2007; but see Susko and Clubb 2008). For instance, Mitchell and Ankeny (2001) found that, in three *H. matronalis* populations in Ohio, “pink” flowers had 6% larger petals than “pale” flowers, and pollinators preferentially visited plants with larger flowers (Weeks and Frey 2007). In contrast, Majetic et al. (2008) did not find any variation in pollinator visitation among *H. matronalis* color morphs in their experimental arrays. To explain the systematic variation in morph frequencies we observed, differences among morphs in pollen fitness would have to vary across the urban-rural gradient, potentially via changes in the composition of pollinator communities or co-flowering species.

Floral pigments can affect plant fitness in a variety of ways, including resistance to herbivory (Simms and Bucher 1996; Fineblum and Rausher 1997; Johnson et al. 2015). For example, anthocyanin, the pigment involved in flower color variation in *H. matronalis* (Majetic 2008), reduces herbivory in several species (Irwin et al. 2003; Strauss et al. 2004; Vaidya et al. 2018). Much of the variation in seed production among the plants we sampled was due to seed predation by the weevil *C. inaffectatus* and we predicted that the purple morph by virtue of higher anthocyanin concentrations would experience lower seed predation. Because this specialist seed predator was first observed in Ontario only a few years before our study (2018, Pentinsaari et al. 2019) we also expected it would be most prevalent in areas with higher human activity. Neither of these expectations were supported by our results. Seed predation was highest on the purple morph and declined with increasing urbanization. This result is not unprecedented. In *Claytonia virginica*, a species with a pink and white flower color polymorphism, plants with more pigmented flowers have lower levels of defensive flavanols and receive more damage from herbivorous slugs (Harborne 1979; Frey 2004). Additionally, some specialist insect herbivores, can be unaffected and even attracted to the defensive compounds of their host (Giamoustaris and Mithen 1995; Lankau 2007; though see Agrawal and Kurashige 2003). Future research should more directly investigate the relation between floral pigmentation and the concentration of defensive secondary metabolites in *H. matronalis*.

The relatively recent arrival of *C. inaffectatus* may have obscured past fitness differences between morphs that are still reflected in the urban-rural cline in color morph frequency variation that we observed. Pre-dispersal seed predators often inflict selection on the same floral display traits as other selective agents such as pollinators, though in opposing directions (Ehrlén et al. 2002; Cariveau et al. 2004; Pérez-Barrales et al. 2013). Our results show that seed predation was highest on purple morphs, especially at the rural end of the gradient. Hence the recent arrival of this seed predator could erase a previous seed fitness advantage of the purple morph in rural populations which could have explained the increasing dominance of the purple morph in rural areas. This is partially supported by our analysis of estimated seed production per plant in the absence of predation. Relative potential seed production of the purple and pink morphs (when the effect of predation was removed) was higher than that of the white morph in rural areas and lower than that of the white morph in urban areas. This “ghost of fitness differences past” hypothesis could be tested by manipulating *C. inaffectatus* seed predation by applying insecticides and by comparing our results to surveys of morph frequencies along replicate urban-rural gradients in regions where *C. inaffectatus* has yet to invade.

## AUTHOR CONTRIBUTIONS

CGE conceived the project and trained the other authors. KGM, with help from CGE, quantified and analyzed color variation among plants. LE and MMK located and sampled the stands of *H. matronalis* and tagged individual plants in 2021. KD and CDM resampled them in 2022 and 2023. In 2021, LJ and MMK harvested tagged plants and LJ measured them. KGM and CGE analyzed the data and wrote the paper with help from all co-authors.

## ACKNOWLEDGEMENTS

The authors thank Spencer C.H. Barrett for long-ago discussions that motivated this study, Rishona Vemulapalli and Ghazal Khonsari for help in the field, Louisa Bartkovich for her help as a sounding board during manuscript revisions, and the Natural Sciences and Engineering Research Council of Canada (NSERC) for funding through the Discovery Grant Program (RGPIN 04831/2020).

## DATA AVAILABILITY STATEMENT

Data will be archived at DRYAD (https://datadryad.org/)

## SUPPORTING INFORMATION

Additional supporting information may be found online in the Supporting Information section at the end of the article.

**Appendix S1.** Locations of study sites and variation in human activity

**Appendix S2.** Spectrometric analysis of flower color variation

**Appendix S3.** Combined analysis of data from 2021, 2022 and 2023

**Appendix S4.** Analysis of the between-year change in color morph diversity

**Appendix S5.** Variation in color morph diversity among generations

**Appendix S6.** Spatial distribution of stands with a high frequency of the purple morph

**Appendix S7.** Retailers of *Hesperis matronalis* seed in North America

