## Supplemental 1 for "Random and non-random variation in flower color along an urban-rural gradient in the introduced mustard *Hesperis matronalis*"

**Appendix S1 – Locations of study sites and variation in human activity**

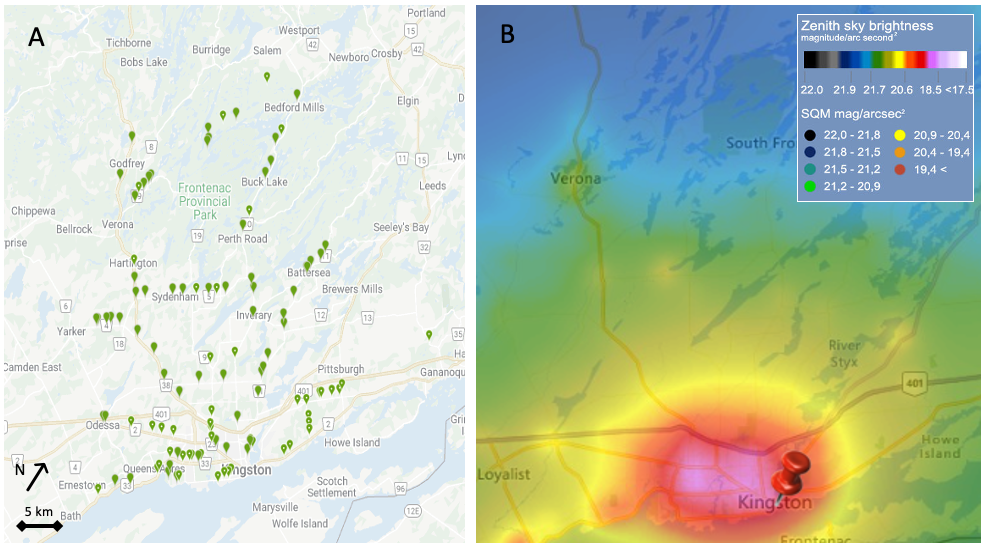

Figure S1. Map of study region in eastern Ontario Canada around the city of Kingston. (A) Green tags indicate the location of each stand of 132 stands of *Hesperis matronalis* surveyed. (B) Map of zenith sky brightness (mag/arcsec^2^, from lightpollutionmap.com) used as a proxy for human activity. Larger zenith sky brightness values (darker blues) indicates lower human activity hence we used “Night Sky Brightness” (NSB = –1x zenith sky brightness) as a measure of human activity.

Table S1. Site codes, coordinates, stand size, and sample size of all stands surveyed and sampled. BP3-26 were not included in the population survey and were only used to collect samples for spectral measurements.

| Site | Latitude | Longitude | NSB | Stand Size (*N*) | | | Sample size (*n*) | | |
| --- | --- | --- | --- | --- | --- | --- | --- | --- | --- |
|  |  |  |  | 2021 | 2022 | 2023 | 2021 | 2022 | 2023 |
| AD-1 | 44.29442 | -76.38790 | -21.18 | 141 | 94 | 58 | 137 | 94 | 63 |
| AD-2 | 44.27802 | -76.38517 | -21.12 | 53 | 30 | 18 | 53 | 30 | 18 |
| AD-3 | 44.27447 | -76.38496 | -21.12 | 250 | 237 | 152 | 53 | 232 | 152 |
| AD-4 | 44.27115 | -76.38444 | -21.12 | 28 | 47 | 37 | 28 | 42 | 37 |
| AD-5 | 44.26436 | -76.38361 | -21.08 | 200 | 167 | 96 | 41 | 166 | 83 |
| BP-2 | 44.24972 | -76.47861 | -19.63 | 200 | NA | NA | 89 | NA | NA |
| BR-10 | 44.45072 | -76.35761 | -21.75 | 27 | 68 | 19 | 27 | 54 | 13 |
| BR-11 | 44.32692 | -76.45870 | -21.04 | 150 | 164 | 111 | 40 | 152 | 93 |
| BR-12 | 44.34093 | -76.45077 | -21.25 | 2 | 0 | 2 | 2 | NA | 2 |
| BR-13 | 44.38152 | -76.42547 | -21.42 | 55 | 81 | 150 | 55 | 76 | 127 |
| BR-14 | 44.42660 | -76.39442 | -21.63 | 10 | 25 | 250 | 10 | 25 | 141 |
| BR-15 | 44.42733 | -76.39261 | -21.63 | 6 | 3 | 46 | 6 | 3 | 46 |
| BR-16 | 44.42840 | -76.39034 | -21.63 | 150 | 31 | 116 | 35 | 24 | 95 |
| BR-17 | 44.42886 | -76.38871 | -21.63 | 6 | NA | 12 | 6 | NA | 12 |
| BR-18 | 44.42918 | -76.38667 | -21.63 | 26 | 53 | 12 | 26 | 53 | 12 |
| BR-2 | 44.30252 | -76.46676 | -20.7 | 54 | 60 | NA | 54 | 32 | NA |
| BR-3 | 44.32261 | -76.46099 | -21.04 | 222 | 400 | 500 | 175 | 83 | 77 |
| BR-6 | 44.37230 | -76.42435 | -21.43 | 69 | 0 | 66 | 69 | NA | 66 |
| BR-7 | 44.40442 | -76.40894 | -21.56 | 188 | 150 | 300 | 188 | 105 | 163 |
| BR-8 | 44.43491 | -76.38155 | -21.63 | 200 | 121 | 200 | 111 | 116 | 177 |
| BR-9 | 44.44219 | -76.36507 | -21.7 | 105 | 44 | 61 | 104 | 44 | 61 |
| CL-1 | 44.56975 | -76.54761 | -21.89 | 11 | 42 | 20 | 11 | 42 | 20 |
| CL-2 | 44.56675 | -76.54818 | -21.89 | 400 | 250 | 1000 | 68 | 141 | 181 |
| CL-3 | 44.56328 | -76.54839 | -21.89 | 62 | 45 | 46 | 62 | 44 | 46 |
| CL-5 | 44.56132 | -76.54728 | -21.88 | 82 | 38 | 15000 | 82 | 38 | 78 |
| CL-6 | 44.55777 | -76.54812 | -21.88 | 83 | 87 | 78 | 83 | 87 | 78 |
| CL-7 | 44.55663 | -76.54892 | -21.88 | 85 | 109 | 127 | 85 | 109 | 127 |
| CR6-2 | 44.27694 | -76.71428 | -20.94 | 141 | 66 | 183 | 141 | 54 | 150 |
| DEV-1 | 44.62120 | -76.45210 | -21.88 | 10 | 0 | 10 | 10 | NA | 10 |
| DEV-2 | 44.58212 | -76.52370 | -21.89 | 18 | 16 | 24 | 18 | 16 | 24 |
| DEV-4 | 44.58504 | -76.50212 | -21.89 | 1 | 0 | 0 | 1 | NA | NA |
| DL-1 | 44.51409 | -76.65222 | -21.83 | 36 | 13 | 26 | 36 | 13 | 26 |
| DL-2 | 44.51017 | -76.66024 | -21.8 | 143 | 162 | 222 | 143 | 162 | 222 |
| DL-3 | 44.52281 | -76.64069 | -21.85 | 4 | 5 | 20 | 4 | 5 | 20 |
| DL-4 | 44.52083 | -76.64423 | -21.84 | 173 | 138 | 500 | 173 | 138 | 175 |
| DL-5 | 44.50090 | -76.66631 | -21.79 | 277 | 500 | 500 | 277 | 222 | 176 |
| DL-6 | 44.49822 | -76.67483 | -21.77 | 50 | 4 | 96 | NA | 4 | 96 |
| FR-1 | 44.21918 | -76.58540 | -20.27 | 200 | 150 | 150 | 96 | 109 | 109 |
| FR-2 | 44.21769 | -76.59366 | -20.34 | 500 | 350 | 129 | 128 | 144 | 109 |
| FR-3 | 44.21981 | -76.58711 | -20.27 | 150 | 250 | 212 | 37 | 162 | 92 |
| FR-4 | 44.21681 | -76.59657 | -20.34 | 2000 | 300 | 300 | 48 | 154 | 174 |
| FR-5 | 44.21511 | -76.60457 | -20.44 | 500 | 2000 | NA | NA | 214 | NA |
| FR-6 | 44.21629 | -76.59883 | -20.34 | 1000 | NA | NA | 50 | NA | NA |
| FR-7 | 44.21657 | -76.59898 | -20.34 | 2000 | 1500 | 1500 | 56 | 246 | 147 |
| FR-8 | 44.21694 | -76.59528 | -20.34 | 500 | 300 | 300 | 66 | 146 | 167 |
| HWY10-1 | 44.27722 | -76.50019 | -19.65 | 92 | 300 | 300 | 92 | 183 | 153 |
| HWY10-10 | 44.48623 | -76.48104 | -21.78 | 1000 | 1000 | 10000 | 74 | 253 | 249 |
| HWY10-11 | 44.55984 | -76.43845 | -21.88 | 76 | 49 | 83 | 76 | 49 | 83 |
| HWY10-12 | 44.56854 | -76.42901 | -21.88 | 76 | 31 | 46 | 76 | 31 | 46 |
| HWY10-14 | 44.52495 | -76.45468 | -21.84 | 93 | 33 | 132 | 93 | 33 | 130 |
| HWY10-17 | 44.53680 | -76.44579 | -21.86 | 57 | 20 | 62 | 57 | 20 | 58 |
| HWY10-18 | 44.60148 | -76.40604 | -21.88 | 200 | 217 | NA | 54 | 216 | NA |
| HWY10-19 | 44.60416 | -76.40325 | -21.88 | 85 | 152 | 500 | 85 | 149 | 378 |
| HWY10-3 | 44.34277 | -76.50422 | -21.16 | 28 | 13 | 250 | 28 | 13 | 158 |
| HWY10-4 | 44.38380 | -76.47490 | -21.4 | 65 | 300 | 15000 | 65 | 176 | 176 |
| HWY10-5 | 44.40812 | -76.47411 | -21.55 | 126 | 300 | 500 | 126 | 253 | 177 |
| HWY10-6 | 44.41337 | -76.47637 | -21.57 | 101 | 100 | 65 | 80 | 83 | 55 |
| HWY10-7 | 44.41749 | -76.47763 | -21.57 | 63 | 50 | 62 | 63 | 51 | 62 |
| HWY10-8 | 44.46621 | -76.49201 | -21.69 | 267 | 88 | 195 | 267 | 87 | 195 |
| HWY10-9 | 44.47117 | -76.49131 | -21.74 | 500 | 500 | 500 | 79 | 168 | 183 |
| HWY2-1 | 44.24426 | -76.4251 | -20.41 | 500 | NA | NA | 204 | NA | NA |
| HWY2-10 | 44.26564 | -76.62484 | -20.36 | 100 | 112 | 800 | 32 | 112 | 223 |
| HWY2-11 | 44.26542 | -76.62315 | -20.36 | 500 | 400 | 500 | 42 | 211 | 163 |
| HWY2-13 | 44.26332 | -76.60724 | -19.99 | 33 | 47 | 2 | 33 | 48 | 2 |
| HWY2-14 | 44.26284 | -76.60439 | -19.99 | 100 | 244 | 250 | 49 | 244 | 294 |
| HWY2-15 | 44.25667 | -76.54202 | -19.31 | 250 | 253 | 2000 | 51 | 189 | 191 |
| HWY2-2 | 44.24778 | -76.41500 | -20.76 | 400 | NA | NA | 217 | NA | NA |
| HWY2-3 | 44.25263 | -76.53531 | -19.29 | 109 | 100 | NA | 108 | NA | NA |
| HWY2-4 | 44.26601 | -76.62498 | -20.36 | 2000 | 2500 | 10000 | 47 | 332 | 295 |
| HWY2-5 | 44.26662 | -76.63214 | -20.49 | 500 | 350 | 500 | 51 | 217 | 196 |
| HWY2-6 | 44.27717 | -76.71808 | -20.9 | 200 | 200 | 111 | 45 | 146 | 101 |
| HWY2-7 | 44.27208 | -76.67288 | -20.97 | 42 | NA | NA | 42 | NA | NA |
| HWY2-9 | 44.26741 | -76.63809 | -20.63 | 13 | 12 | 250 | 13 | 12 | 159 |
| HWY33-1 | 44.23789 | -76.55895 | -19.17 | 2000 | 250 | 1000 | 123 | 168 | 255 |
| HWY33-10 | 44.23683 | -76.56381 | -19.47 | 47 | 4 | 14 | 47 | 4 | 14 |
| HWY33-12 | 44.23081 | -76.62550 | -20.45 | 30 | 13 | 52 | NA | 13 | 52 |
| HWY33-13 | 44.22784 | -76.62732 | -20.47 | 200 | 80 | 15 | 27 | 60 | 8 |
| HWY33-14 | 44.21383 | -76.67445 | -20.76 | 161 | 100 | 200 | 161 | 82 | 124 |
| HWY33-16 | 44.21180 | -76.70017 | -21.06 | 24 | 250 | 166 | 24 | 240 | 60 |
| HWY33-17 | 44.21228 | -76.69859 | -21.09 | 20 | 3 | 5 | 20 | 3 | 3 |
| HWY33-2 | 44.23780 | -76.57447 | -19.15 | 67 | 4 | 4 | 62 | 4 | 4 |
| HWY33-3 | 44.23784 | -76.57909 | -19.32 | 148 | NA | NA | 126 | NA | NA |
| HWY33-4 | 44.23743 | -76.58957 | -19.74 | 500 | 200 | 400 | 76 | 167 | 184 |
| HWY33-5 | 44.24023 | -76.59729 | -19.72 | 239 | NA | NA | 222 | NA | NA |
| HWY33-6 | 44.24111 | -76.60995 | -20.1 | 216 | 200 | 350 | 168 | 118 | 238 |
| HWY33-7 | 44.22481 | -76.62972 | -20.47 | 5000 | 2000 | 1000 | 73 | NA | NA |
| HWY33-8 | 44.20267 | -76.72618 | -21.01 | 300 | 300 | 500 | 63 | 151 | 142 |
| HWY38-1 | 44.30212 | -76.59492 | -20.66 | 52 | 167 | 26 | 52 | 167 | 26 |
| HWY38-11 | 44.43664 | -76.66788 | -21.61 | 24 | 13 | 19 | 24 | 13 | 19 |
| HWY38-12 | 44.41811 | -76.66705 | -21.53 | 12 | 119 | 300 | 12 | 119 | 300 |
| HWY38-14 | 44.40346 | -76.66501 | -21.47 | 183 | 200 | 400 | 183 | 199 | 308 |
| HWY38-2 | 44.32018 | -76.61892 | -20.66 | 1000 | 1000 | 1000 | 192 | 200 | 192 |
| HWY38-3 | 44.34702 | -76.63527 | -21.35 | 63 | 38 | 72 | 63 | 38 | 72 |
| HWY38-4 | 44.36467 | -76.66261 | -21.46 | 250 | 600 | 600 | 84 | 305 | 195 |
| HWY38-5 | 44.52327 | -76.69119 | -21.79 | 200 | 100 | 500 | 53 | 57 | 162 |
| HWY38-6 | 44.56074 | -76.67168 | -21.88 | 31 | 0 | 19 | 31 | NA | 19 |
| JR-1 | 44.30887 | -76.33063 | -21.39 | 53 | 50 | 58 | 43 | 47 | 59 |
| LOP-1 | 44.21625 | -76.53180 | -19.82 | 179 | 102 | 75 | 179 | 102 | 75 |
| LOP-2 | 44.21921 | -76.53400 | -19.89 | 49 | 22 | 86 | 49 | 22 | 85 |
| LP-1 | 44.22134 | -76.60639 | -20.28 | 108 | 85 | 50 | 108 | 85 | 50 |
| LP-2 | 44.22049 | -76.61205 | -20.55 | 157 | 95 | 77 | 157 | 93 | 71 |
| LP-3 | 44.22686 | -76.60985 | -20.41 | 71 | 78 | 70 | 71 | 78 | 68 |
| MR-1 | 44.29397 | -76.40295 | -21.03 | 208 | 29 | 25 | 200 | 28 | 22 |
| MR-2 | 44.30009 | -76.38939 | -21.21 | 10000 | 2000 | 2000 | 82 | 240 | 185 |
| MR-3 | 44.30135 | -76.37460 | -21.31 | 4 | 0 | NA | 4 | NA | NA |
| MR-4 | 44.30152 | -76.37225 | -21.31 | 150 | 79 | 37 | 35 | 74 | 37 |
| MR-5 | 44.30180 | -76.36663 | -21.34 | 24 | 6 | 14 | 24 | 6 | 14 |
| MR-6 | 44.30190 | -76.36472 | -21.34 | 250 | 250 | 200 | 57 | 174 | 203 |
| MR-7 | 44.30367 | -76.34739 | -21.38 | 4 | 0 | NA | 4 | NA | NA |
| MR-8 | 44.30472 | -76.33424 | -21.37 | 500 | 500 | 750 | 77 | 178 | 136 |
| POH-1 | 44.22077 | -76.51494 | -19.88 | 6 | 0 | 0 | 6 | NA | NA |
| PV-1 | 44.22045 | -76.52095 | -19.82 | 92 | 140 | 88 | 92 | 141 | 83 |
| RR-10 | 44.40682 | -76.60550 | -21.33 | 6 | 56 | 52 | 6 | 56 | 42 |
| RR-12 | 44.40515 | -76.65005 | -21.51 | 52 | 19 | 250 | 52 | 20 | 250 |
| RR-3 | 44.40978 | -76.51814 | -21.56 | 500 | 1500 | 10000 | 98 | 342 | 318 |
| RR-4 | 44.40735 | -76.53360 | -21.55 | 69 | 160 | 300 | 69 | 161 | 173 |
| RR-5 | 44.40747 | -76.56728 | -21.5 | 300 | 400 | 500 | 73 | 174 | 234 |
| RR-7 | 44.40912 | -76.51962 | -21.56 | 124 | 500 | NA | 124 | 271 | NA |
| RR-8 | 44.40723 | -76.54723 | -21.54 | 29 | NA | 119 | 29 | NA | 112 |
| RR-9 | 44.40717 | -76.58812 | -21.39 | 250 | 250 | 100 | 57 | 173 | 67 |
| SJ-1 | 44.24558 | -76.51776 | -19.15 | 300 | 209 | 100 | 45 | 165 | 36 |
| SJ-4 | 44.27089 | -76.51781 | -19.34 | 300 | 1500 | NA | NA | 256 | NA |
| SR-1 | 44.25980 | -76.54376 | -19.31 | 10000 | 500 | 10000 | 102 | 168 | 154 |
| SR-2 | 44.26597 | -76.54379 | -19.49 | 500 | 500 | 10000 | 57 | 276 | 177 |
| SR-3 | 44.26987 | -76.54380 | -19.49 | 171 | 35 | 2000 | 168 | 28 | 265 |
| SR-4 | 44.28353 | -76.54200 | -19.98 | 423 | 15 | 250 | 423 | 15 | 229 |
| SR-5 | 44.33706 | -76.54413 | -21.06 | 113 | NA | NA | 113 | NA | NA |
| UR-1 | 44.31953 | -76.52275 | -20.81 | 123 | 300 | 1000 | 123 | 192 | 196 |
| UR-2 | 44.31835 | -76.56247 | -20.81 | 23 | 0 | 0 | 23 | NA | NA |
| YR-1 | 44.37771 | -76.69018 | -21.53 | 12 | 0 | 32 | 12 | NA | 32 |
| YR-2 | 44.37773 | -76.70595 | -21.57 | 200 | 500 | 1000 | 70 | 161 | 299 |
| YR-3 | 44.37768 | -76.71088 | -21.58 | 97 | 151 | 140 | 97 | 151 | 134 |
| YR-4 | 44.37701 | -76.72179 | -21.59 | 215 | 280 | 350 | 215 | 277 | 319 |
| YR-5 | 44.37762 | -76.71265 | -21.58 | 74 | 253 | 250 | 74 | 258 | 238 |
| YR-6 | 44.37650 | -76.72721 | -21.6 | 26 | 0 | 5 | 26 | NA | 5 |
| YR-7 | 44.37641 | -76.72894 | -21.6 | 11 | 0 | 4 | 11 | NA | 4 |
