## Supplemental 2 for "Random and non-random variation in flower color along an urban-rural gradient in the introduced mustard *Hesperis matronalis*"

**Appendix S2 – Spectrometric analysis of flower colour variation**

Based on visual inspection of thousands of flowers across many stands, we felt we could reliably distinguish three discrete flower colour morphs: white (W), pink (P) and purple (V ~ violet). A set of standard photographs were used during population surveys to ensure consistency in colour morph scoring (Fig. 1). To quantify flower colour variation in *H. matronalis* more objectively and to determine whether this 3-morph classification adequately captured flower colour variation, we collected 988 flowers (one flower per plant) from four sites: ALW (48 flowers), BP (580), Hwy2-4 (200), and MR-2 (160) (Table S1). Individual flowers were pinched off from the inflorescence, leaving ≥ 1.5 cm of pedicel. To control for potential changes in flower colour with age, all flowers were collected when the stamens were just beginning to emerge from the flower tube. We transported flowers at 4°C to the lab, where they were stored at 4°C and high humidity until the colour was measured (within 5 h) with a spectrometer.

Spectrometry

We measured colour with a USB4000 spectrometer with a PX-2 pulsed xenon lamp set to single flash (Ocean Optics, Inc. Dunedin, FL) OceanView 2.0.8 software measured reflectance across from 300–700 nm with integration time set to 100 ms and boxcar width to 5. Reflectance values outside this range were error prone. We made three replicate measures per flower involving three different petals. A spectralon white standard and a black dark standard were used to update the reference light and background spectra, respectively, before each flower was measured.

Analysis of Spectral Data

Spectral data was analyzed using the pavo R package (version 1.43.17; Maia et al. 2019). A single spectrum was calculated per flower from the average of the three petal measurements and the data were smoothed by a span of 0.2. For flowers classified as white reflectance steadily increased from 350–450 nm (Fig. S2), peaking just beyond 450 nm and remaining high to 700 nm. In contrast, flowers classified as purple showed two reflectance peaks at 450 nm and 690 nm and a low point at 560 nm. The peak at 690 nm was almost always higher than the 450 nm peak. Pink flowers exhibited more variation in the spectrum shape between white and purple. The relative percent of reflectance at 560 nm captured the primary axis of flower colour variation, so we calculated a “spectral shape” parameter following Frey (2004) as (reflectance at 450 nm + reflectance 690 nm)/2 – reflectance at 560 nm). This value increases as the depth of the spectral “valley” increased such that pure white flowers have a value of zero. Spectral shape varied widely among flowers (Fig. S2) and was distributed with two modes and much variation in between (Fig. S3). To determine how much variation in spectral shape was accounted for by colour morph, we calculated the coefficient of determination (*r*^2^) from a linear model with morph as a predictor fit to variation in spectral shape. More than 70% of the variation in spectral shape was distributed among the three flower colour morphs distinguished in this study (Fig. S4; *r*^2^ = 0.737, *F*_2,985_ = 1380, *P* < 0.0001). Neither spectral shape or the pattern of variation in spectral shape among morphs varied among stands (among stands: *F*_3,982_ = 0.77, *P* = 0.51; interaction between morph and stand *F*_6,976_ = 1.04, *P* = 0.40).

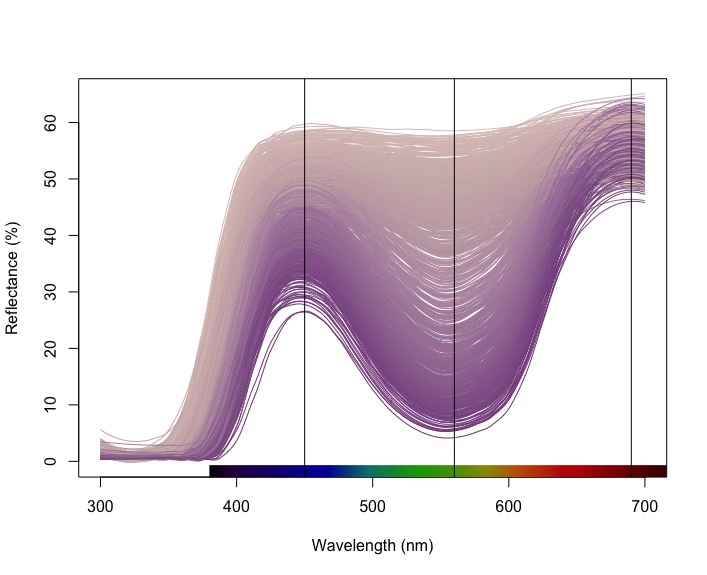

Figure S2. Variation in reflectance between 300–700 nm among 988 flowers of *Hesperis matronalis* flowers sampled from four stands in eastern Ontario, Canada. Each spectrum is for a single flower averaged from spectral measurements of three petals per flower. Line colour is an approximation of the perceived colour and was converted from the reflectance spectra using the spec2rgb() function in the pavo R package. The black horizontal lines at 450, 560, and 690 nm mark the peaks and valley of the spectra used to calculate spectral shape.

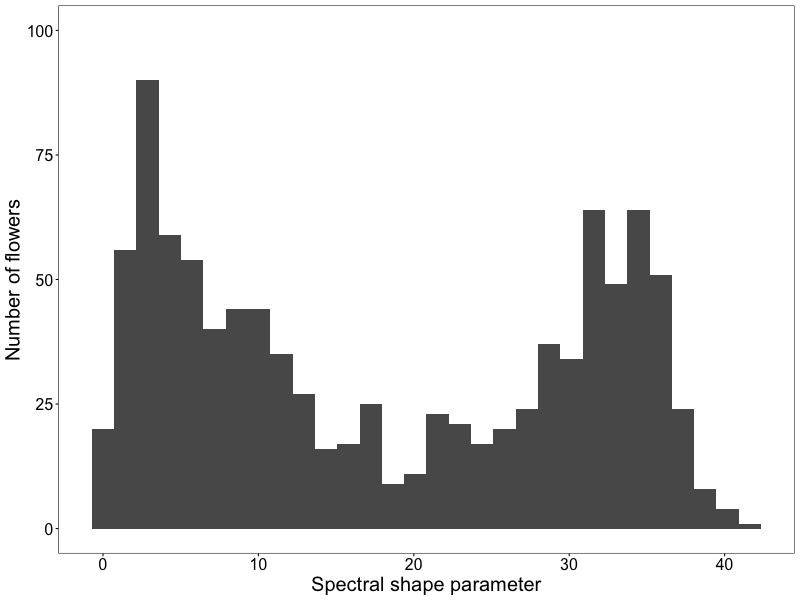

Figure S3. Variation in spectral shape parameter between 988 flowers of *Hesperis matronalis* sampled from four stands in eastern Ontario, Canada. The spectral shape parameter is the difference between the peaks and the low point of the spectral distribution of each flower and was calculated as: (reflectance at 450 nm + reflectance at 690 nm)/2 – reflectance at 560 nm (see Fig S2). Light coloured flowers have a low value of spectral shape and darker purple flowers have higher values.

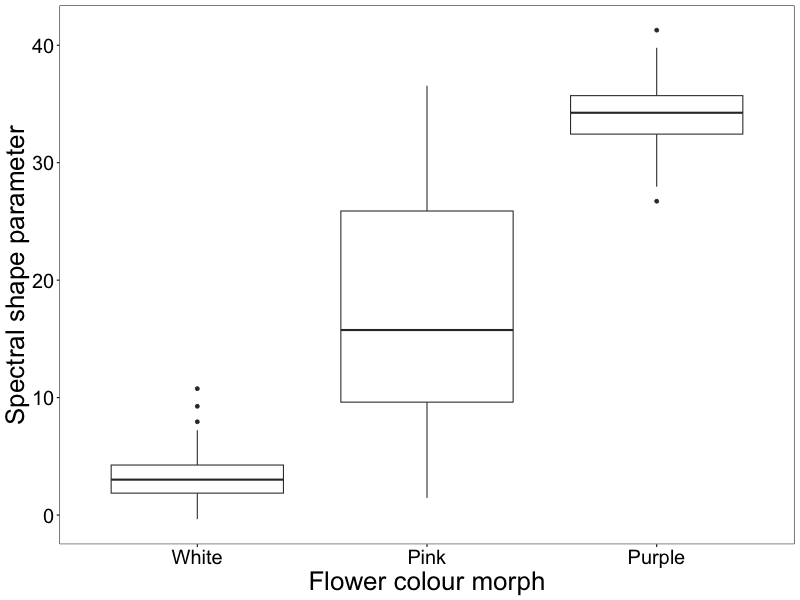

Figure S4. Variation in the spectral shape parameter among three discrete flower colour morphs observed in *Hesperis matronalis* among four stands in eastern Ontario, Canada. Data are from 246 white, 498 pink and 244 purple flowers. 73% of variation in spectral shape is distributed among rather within colour morphs. The thick middle line in each box plot indicates the median, the top and bottom of the box indicate the upper and lower quartile, and whiskers extend to the maximum and minimum points or 1.5 interquartile range from the median whatever is less.
