## Supplemental 3 for "Random and non-random variation in flower color along an urban-rural gradient in the introduced mustard *Hesperis matronalis*"

**Appendix S3 – Combined analysis of data from 2021, 2022 and 2023**

Table S2. Analyses of variation in colour morph diversity (*H*’) among stands of *Hesperis matronalis.* This analysis involves the 105 stands that were sampled in all three years.

The colour morph diversity and size of individual stands correlated positively between years (*H*’: Kendall’s *W* = +0.75, *P* < 0.0001; log_10_*N*: *W* = +0.80, *P* < 0.0001;). Variation in colour morph diversity was fit to mixed-effects linear models with human activity (NSB), stand size (log_10_*N*) and year as fixed effects as well as 2-way interactions between year and both stand size and human activity. A 3-way interaction was not included because the interaction between human activity and stand size was not significant in either year (Table 1). Stand was included as a random effect. Models were fit using the *lmer* function in the lme4 R package (version 1.1-33, https://CRAN.R-project.org/package=lme4). Significance of fixed effects were evaluated using likelihood-ratio tests. Cells include tests of significance (χ^2^, *P*) and standardized partial regression coefficients (*b*) when significant. The significant effect of year results from higher *H*’ in 2023 than 2021 or 2022 (Table S3). Effects of both NSB and log_10_*N* varied slightly among years, with a slightly stronger effect of NSB on *H*’ in 2021 (b = +0.353) than 2023 (*b* = +0.220) and a slightly stronger effect of log_10_*N* on *H*’ in 2022 (*b* = +0.383) than 2023 (*b* = +0.212).

| Predictor |  |
| --- | --- |
| Human activity (NSB) | *b* = +0.367  χ^2^ = 14.0, *P* = 0.00018 |
| Stand size (log_10_*N*) | *b* = +0.245  χ^2^ = 19.6, *P* < 0.0001 |
| Year (Y) | *H*’_2023_ > (*H*’_2021_+ *H*’_2022_)  χ ^2^ = 4.1, *P* = 0.042 |
| Y x NSB | *b*_2021_ > *b*_2023_  χ^2^ = 5.7, *P* = 0.058 |
| Y x log_10_*N* | *b*_2022_ > *b*_2023_  χ^2^ = 6.2, *P* = 0.044 |
