## Supplemental 4 for "Random and non-random variation in flower color along an urban-rural gradient in the introduced mustard *Hesperis matronalis*"

**Appendix S4 – Analysis of the between-year change in colour morph diversity**

We evaluated between-year variation in colour morph diversity (*H*’) and log_10_*N* by fitting variation in these parameters to a mixed-effects model with year as a fixed effect and stand as a random effect using the *lmer* function in the lme4 R package. The significance of year was evaluated using a likelihood ratio test. We contrasted the means for each year predicted by the models using the Tukey method implemented by the *pairs* function in the emmeans R package (version 1.8.7, https://CRAN.R-project.org/package=emmeans). Both *H*’ and log_10_*N* were higher in 2023 than in the two previous years (Table S3).

We investigated whether year-to-year variation in *H*’ among sites was associated with the degree of urbanization, the within-site change in stand size and/or the within-site variation in sample size by fitting the difference in *H*’ between 2023 and the average of 2021 and 2022 (Δ*H*’ = *H*’_2023_ – [*H*’_2021_ + *H*’_2022_]/2), as a response variable, to a linear model with NSB, the difference in log_10_*N* (Δlog_10_*N* = log_10_*N*_2023_ – [log_10_*N*_2021_ + log_10_*N*_2022_]/2) and the difference in log_10_-transformed sample size (*n*) as potential predictors with significance of each predictor evaluated using likelihood ratio tests (LRT). The between-years increase in *H*’ was lower in urban than rural stands (standardized partial regression coefficient for NSB, *b* = –0.19 ± 0.095, LRT *F*_1,101_ = 3.99, *P* = 0.048), increased with Δlog_10_*N* though not significantly (*b* = +0.23 ± 0.12, *F*_1,101_ = 3.28, *P* = 0.073) but did not vary with the change in log-transformed sample size (*b* = +0.065 ± 0.12, *F*_1,101_ = 0.27, *P* = 0.60).

Table S3. Variation in colour morph diversity (*H*’) and stand size (log_10_*N*) among years for the 105 eastern Ontario stands of *Hesperis matronalis* sampled in all three years. Each parameter varied significantly among years (*H*’ LRT χ^2^ = 25.69, df = 2, *P* < 0.0001; log10N χ^2^ = 20.43, df = 2, *P* < 0.0001). Cells contain means ± 1SE predicted by the mixed models with superscript letters showing the results of Tukey multiple contrasts. Values not sharing a letter are significantly different.

| Year | *H*’ | log_10_*N* |
| --- | --- | --- |
| 2021 | 0.734 ± 0.031^B^ | 2.11 ± 0.065^B^ |
| 2022 | 0.768 ± 0.031^B^ | 2.02 ± 0.065^B^ |
| 2023 | 0.847 ± 0.031^A^ | 2.26 ± 0.065^A^ |
