## Supplemental 5 for "Random and non-random variation in flower color along an urban-rural gradient in the introduced mustard *Hesperis matronalis*"

**Appendix S5 – Variation in colour morph diversity among generations**

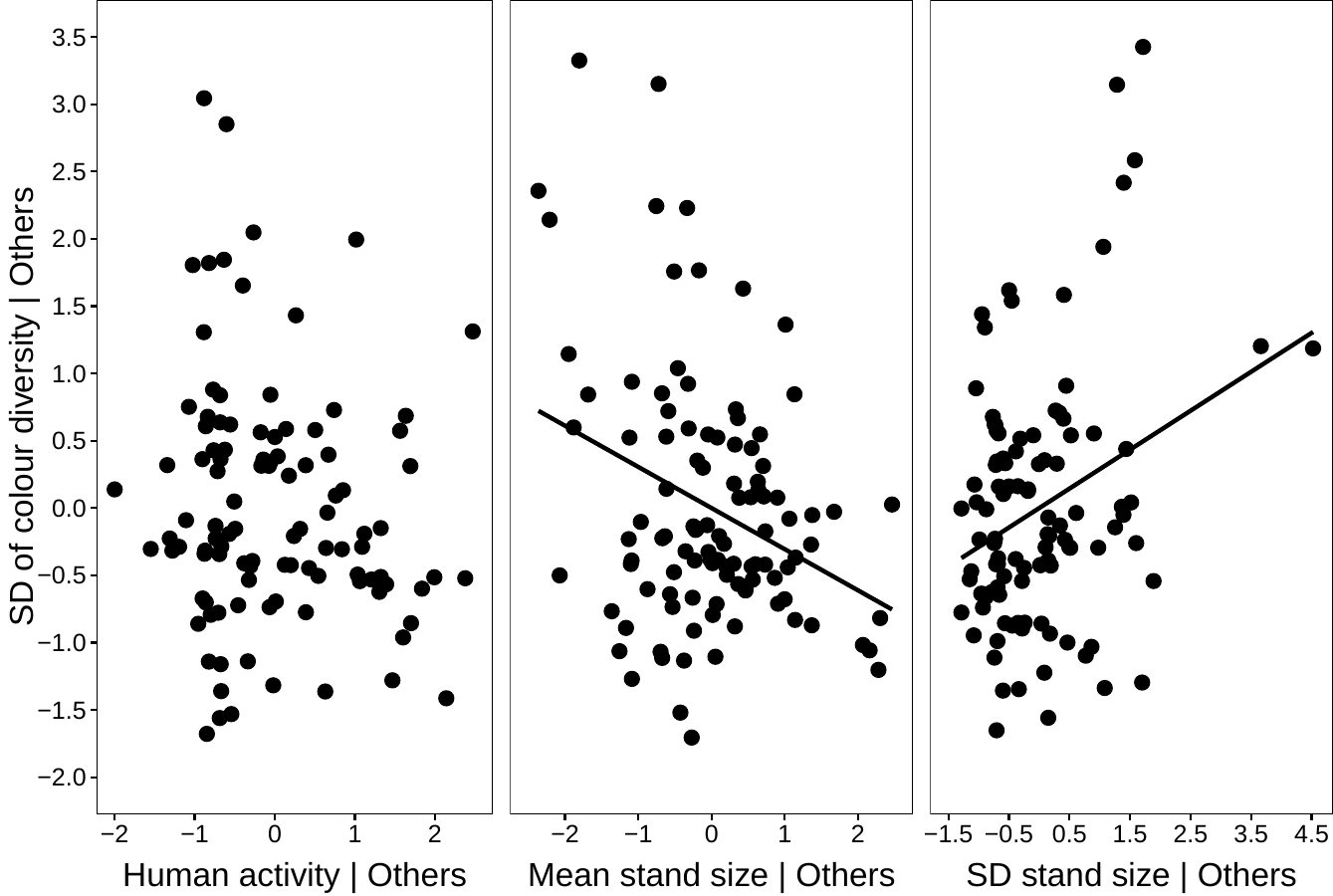

Figure S5. Added-variable plots showing the relations between the standard deviation (SD) of flower colour morph diversity (*H’*) among generations and human activity (NSB) and the mean and standard deviation of stand size (log_10_*N*) across generations in *Hesperis matronalis* from eastern Ontario, Canada (105 stands sampled in all three years). Each point is a stand. The y-axis is residual SD of *H’* when the other two predictors are held constant. The x-axis is the residual of the focal predictor when the other two predictors are held constant.
