## Supplemental 6 for "Random and non-random variation in flower color along an urban-rural gradient in the introduced mustard *Hesperis matronalis*"

**Appendix S6 – Spatial distribution of stands with a high frequency of the purple morph**

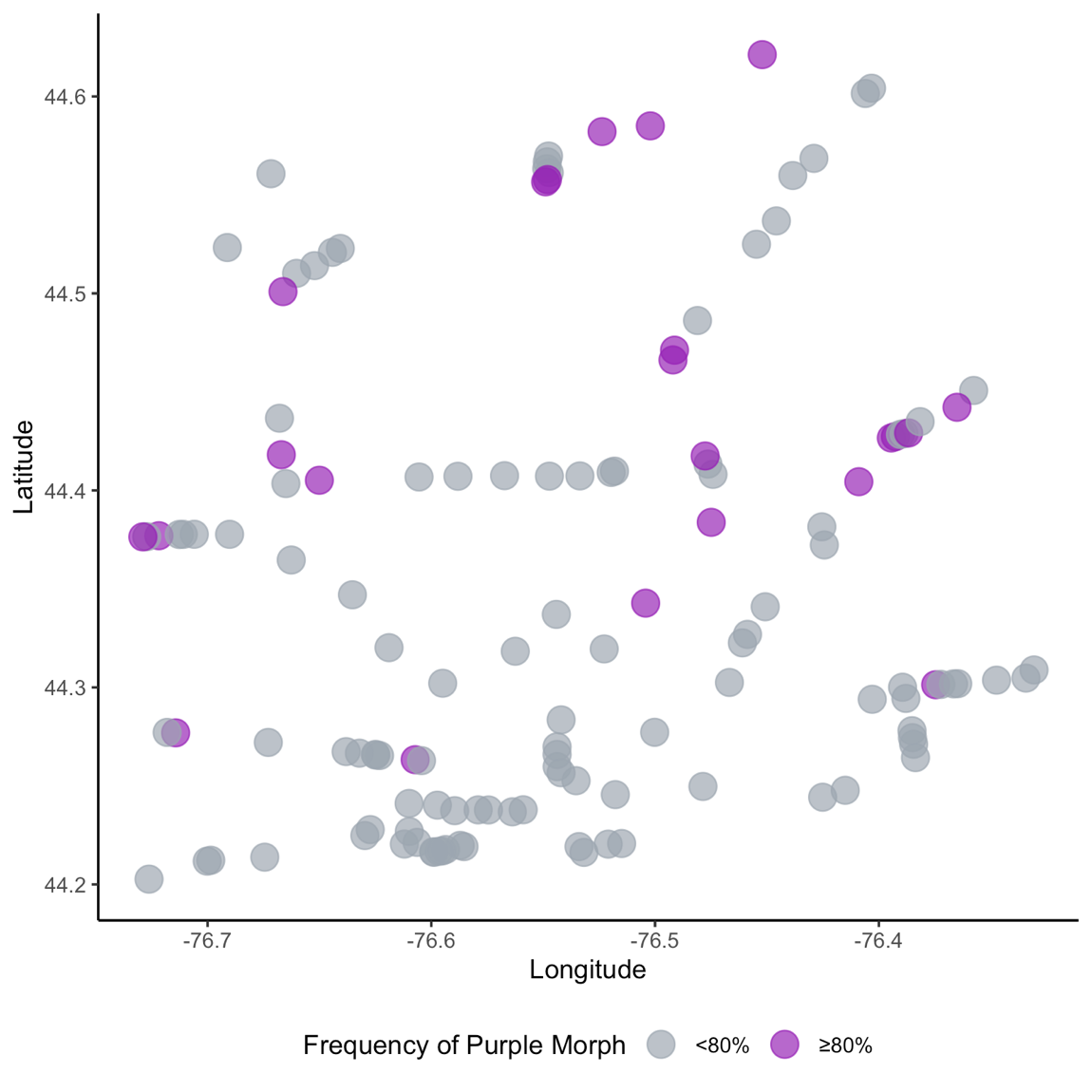

Figure S6. Location of populations surveyed in 2021, highlighting the spatial spread of populations with a high frequency of the purple morph (n = 132). Purple dots indicate populations where over 80% of the plants surveyed were identified as purple morphs. Grey dots indicate populations where less than 80% of the plants surveyed were identified as purple morphs.
