## Supplemental 7 for "Random and non-random variation in flower color along an urban-rural gradient in the introduced mustard *Hesperis matronalis*"

**Appendix S7 – Retailers of *Hesperis matronalis* seed in North America**

In July 2023, we comprehensively searched the internet for North American seed retailers that sold *Hesperis matronalis*. We scored a seed stock as polymorphic for flower colour if it was described as such or if the picture accompanying the product description included more than one colour morph (picture-based scores in brackets below). We scored the seed stock as monomorphic for flower colour if the product description indicated only one colour or if the accompanying photograph included only a single morph (in brackets). For monomorphic seed stock we recorded the colour described or included in the picture (in brackets). Some online retailers did not disclose their location on their website.

| Retailer | Location | Prov/State | Country | Polymorphic? | Monomorphic for | Website |
| --- | --- | --- | --- | --- | --- | --- |
| Wildwood outdoor living center | Victoria | BC | CAN | No | pink | wildwood.express.com |
| Seed Corner | Bellevue | WA | USA | Yes | – | seedcorner.com |
| Sheffield’s Seeds | Locke | NY | USA | Yes | – | sheffields.com |
| Victory Seed Company | Irvine | TX | USA | NA | – | victoryseeds.com |
| Eden Brothers | Arden | NC | USA | Yes | white or purple | www.edenbrothers.com |
| Bumbleseeds | Vancouver Island | BC | CAN | No | purple | bumbleseeds.com |
| Select Seeds | Union | CT | USA | No | white or purple | www.selectseeds.com |
| Everwilde Farms | Fallbrook | CA | USA | No | pink | www.everwilde.com |
| Project Purity Seeds | Metropolis | IL | USA | No | lavender | www.projectpurityseeds.com |
| Ernst Conservation Seeds | Meadville | PA | USA | Yes | – | www.ernstseed.com |
| Outside Pride Seeds | Salem | OR | USA | (Yes) | – | www.outsidepride.com |
| Asho Gardens | – | – | CAN | (Yes) | – | ashogardens.com |
| Heritage Harvest Seed | Fisher Branch | MB | CAN | No | purple | heritageharvestseed.com |
| MySeeds.co | Oakland Gardens | NY | USA | Yes | – | myseeds.co |
| American Meadows | – | – | USA | (Yes) | – | www.americanmeadows.com |
| Metchosin Farm | Metchosin | BC | CAN | No | pink | metchosinfarm.ca |
| Brother Nature Seeds | Victoria | BC | CAN | (Yes) | – | brothernature.ca |
| Swallowtail Garden Seeds | Santa Rosa | CA | USA | (Yes) | – | www.swallowtailgardenseeds.com |
| Ferri Seeds | – | ON | CAN | (No) | (purple) | ferriseeds.com |
| Idoville315 (eBay) | – | NA | NA | No | purple | www.ebay.ca |
| Prairie Garden Seeds | North Battledford | SK | CAN | Yes | – | prairiegardenseeds.ca |
| Seedville USA | Canton | OH | CAN | No | purple | seedvilleusa.com |
| Diane’s Flower Seeds | Provo | UT | USA | Yes | – | www.dianeseeds.com |
| Caribbean Garden Seeds | Pottstown | PA | USA | (No) | (purple) | caribbeangardenseed.com |
| Florabunda Seeds | Keene | ON | CAN | (No) | (purple) | www.florabundaseeds.com |
| Wildseed Farm | Fredericksburg | TX | USA | (No) | (purple) | www.wildseedfarms.com |
